# scConfluence : single-cell diagonal integration with regularized Inverse Optimal Transport on weakly connected features

**DOI:** 10.1101/2024.02.26.582051

**Authors:** Jules Samaran, Gabriel Peyré, Laura Cantini

## Abstract

The abundance of unpaired multimodal single-cell data has motivated a growing body of research into the development of diagonal integration methods. However, the state-of-the-art suffers from the loss of biological information due to feature conversion and struggles with modality-specific populations. To overcome these crucial limitations, we here introduced scConfluence, a novel method for single-cell diagonal integration. scConfluence combines uncoupled autoencoders on the complete set of features with regularized Inverse Optimal Transport on weakly connected features. We extensively benchmark scConfluence in several single-cell integration scenarios proving that it outperforms the state-of-the-art. We then demonstrate the biological relevance of scConfluence in three applications. We predict spatial patterns for *Scgn, Synpr* and *Olah* in scRNA-smFISH integration. We improve the classification of B cells and Monocytes in highly heterogeneous scRNA-scATAC-CyTOF integration. Finally, we reveal the joint contribution of *Fezf2* and apical dendrite morphology in Intra Telencephalic neurons, based on morphological images and scRNA.

## Introduction

In the last decade, single-cell transcriptomics (scRNA) has revolutionized our understanding of the diversity of cells constituting living tissues^1–3^. Since then, a new milestone has been reached with the introduction of high-throughput sequencing technologies allowing to measure additional molecular modalities, such as chromatin accessibility (scATAC)^4,5^ and methylation (snmC)^6^, at the resolution of the single cell. More recently, technologies allowing the joint measurement of different single-cell modalities from the same cell (i.e. paired data) have been proposed^7–15^. Examples of these cutting-edge sequencing technologies are CITE-seq, simultaneously measuring RNA and surface protein abundance by leveraging oligonucleotide-conjugated antibodies^8^, and 10x Genomics Multiome platform, quantifying RNA and chromatin accessibility by microdroplet-based isolation of single nuclei.

Different single-cell modalities describe complementary facets of the cell; their joint analysis is thus expected to provide tremendous power to uncover cellular identities^16^. For achieving this aim, paired single-cell multimodal data represent an ideal resource^17,18^ and numerous methods have been designed for their integration^19–22^. Nevertheless, paired data are still rare and limited in the amount of modalities that they contain (maximum three)^23^. Single-cell multimodal data profiled from different cells of the same biological condition, i.e. unpaired data, thus represent a precious resource for accessing different molecular facets of a cell and better understanding its identity.

The integration of unpaired single-cell multimodal data, i.e. diagonal integration, is more challenging than paired integration^24^. Indeed, comparing cells from different modalities is not straightforward, as they are described by different features (e.g. genes, peaks, proteins). The aim of diagonal integration is to define a low dimensional latent space shared by all modalities. In this shared latent space, cells should be arranged according to their biological similarity, independently from their modality of origin. Providing such a biologically meaningful modality alignment of cells, different from the many potential artificial alignments which overlap cells from different cell types, is extremely challenging.

To guide cell alignment between modalities in the shared latent space, diagonal integration leverages prior biological information^24^. Indeed, connections between the features of different modalities are generally known in biology. For instance, chromatin peaks can be mapped to genes based on their proximity to gene promoter regions, thus enabling the computation of gene activity measurements^25,26^. Similarly, protein-coding genes and their corresponding proteins can be used as connections between scRNA-seq and proteomic data. Most of the state-of-the-art methods use this prior biological knowledge to convert all modalities to the same features and then handle the alignment similarly to batch effect correction^27–29^. However, this conversion can result in an important loss of biological information as features across modalities are weakly connected. Indeed, across-modality feature connections are often rare and noisy. For example, protein-coding genes are a subset of all the expressed genes and not all possible chromatin peaks are close to the promoter of a gene. This problem becomes even more challenging once the features measured in one modality are few due to technological limitations (e.g. targeted CyTOF providing only few proteins quantified across cells). State-of-the-art methods not requiring modality conversions also exist^30,31^. However they still depend on the assumption that most features can be reliably connected across modalities. In addition, many state-of-the-art methods^27,28,30,31^ ignore the possibility that a population of cells (cell type/state) can be present only in one modality, which is frequently the case for unpaired data.

Here, we propose scConfluence, a novel diagonal integration method combining uncoupled autoencoders, which reduce the dimensionality of the original data to a shared latent space and account for potential batch effects, together with regularized Inverse Optimal Transport (rIOT)^32^, which aligns cells across modalities in the shared latent space by leveraging weakly connected features. By employing rIOT to ensure modality alignment, scConfluence can independently process the complete set of original features through autoencoders while utilizing only the connected features for aligning cell embeddings. Therefore, our approach does not suffer from the loss of biological information generally resulting from modality conversion prior to dimension reduction. In addition, thanks to the unbalanced relaxation of Optimal Transport^33^, scConfluence can also deal with cell types absent in a modality thus overcoming all the major limitations of the state-of-the-art.

We extensively benchmark scConfluence with respect to the state-of-the-art in several scRNA-surface protein and scRNA-scATAC integration problems. This in-depth comparison proves that scConfluence’s embeddings outperform the state-of-the-art across a wide variety of datasets. We further demonstrate scConfluence’s robustness, accuracy and general applicability in addressing three diverse and crucial biological questions. First, we integrate scRNA-seq and smFISH profiled from mouse somatosensory cortex and predict *Scgn, Synpr* and *Olah* to have spatial patterns of expression amenable for further biological investigation. Second, scConfluence’s integration of scRNA-seq, scATAC-seq and CyTOF improves the classification of B cells and Monocytes in highly heterogeneous human PBMC datasets. Finally, scConfluence integrates neuronal morphological images with scRNA-seq from the mouse primary motor cortex revealing the joint contribution of the Transcription Factor *Fezf2* and apical dendrite morphology to information processing in Intra Telencephalic neurons.

scConfluence is highly modular, allowing its generalization to the new integration scenarios that will arise in consequence of the continuous single-cell technological developments (e.g. single-cell metabolomics). scConfluence is implemented as an extensively documented open-source Python package seamlessly integrated within the scverse ecosystem^34^ and is available at https://github.com/cantinilab/scconfluence.

## Results

### scConfluence a new method for diagonal single-cell multimodal integration

We developed scConfluence, a novel method for single-cell diagonal integration combining uncoupled autoencoders with regularized Inverse Optimal Transport (rIOT) on weakly connected features.

As shown in Figure 1A, the inputs of scConfluence are single-cell data from *M* modalities represented by the matrices 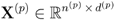 with *p* ∈ [1.. *M* ], where rows correspond to cells and columns to features (e.g. genes, chromatin peaks, proteins). The cells of **X**^(*p*)^ can come from multiple experimental batches. As discussed in the Introduction, although each modality is grounded in a different feature space, across-modality connections between some features can be defined based on prior biological knowledge. Therefore, we expect that for all pairs of modalities (*p, p*′), we have access to 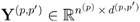 and 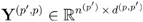, conversions of **X**^(*p*)^ and **X**^**(*p***′)^ to common features, respectively. For example, if *p* corresponds to scRNA and *p*′ is scATAC, **Y**^(*p, p*′)^ and **Y**^(*p*′, *p*)^ correspond to the RNA count matrix and the gene activity matrix derived from peak accessibility counts, respectively.

**Figure 1.**
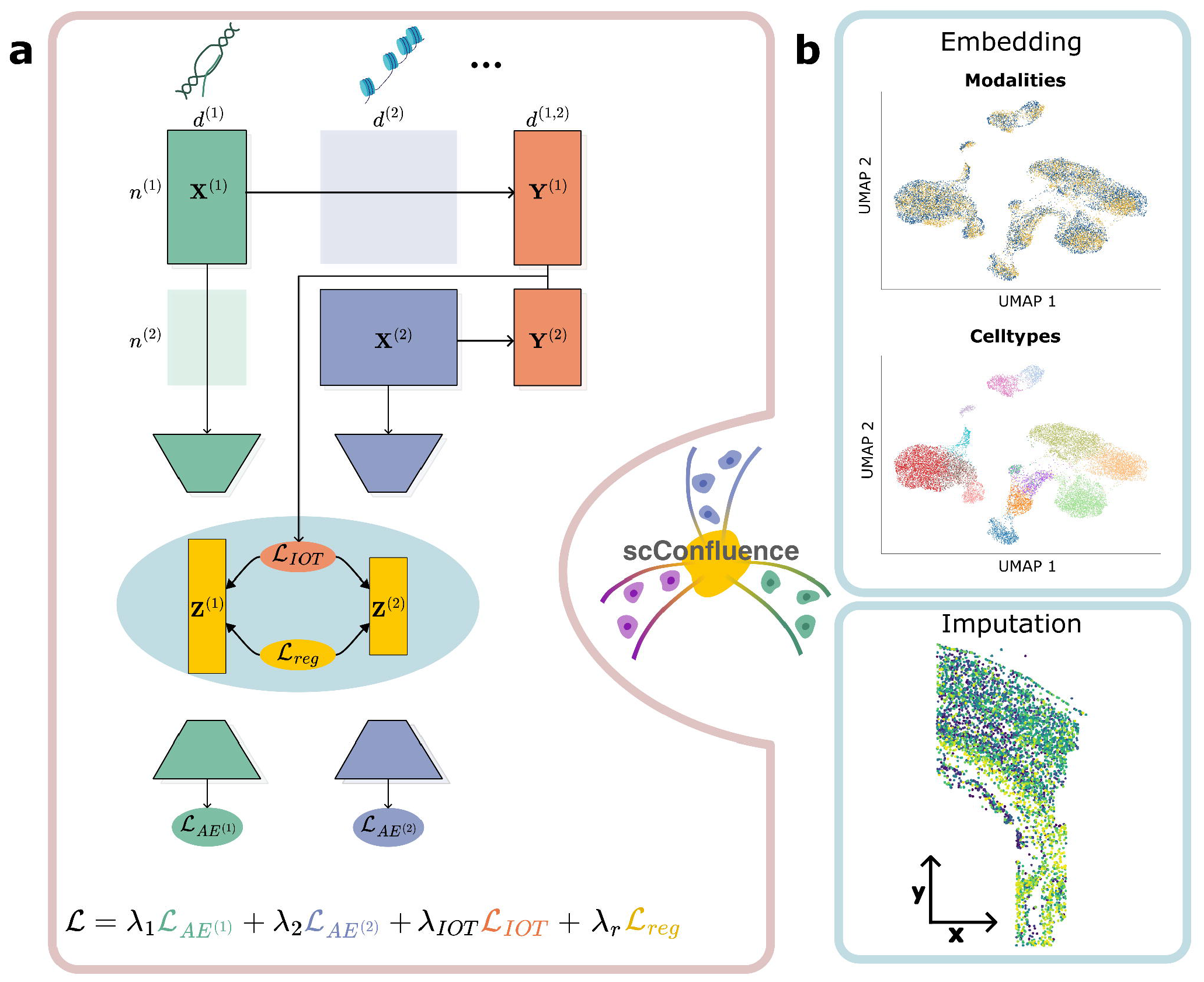
The scConfluence framework for diagonal integration. (a) Schematic representation of the framework simplified to only two modalities (*M* = 2). While the original data matrices **X**^(1)^ and **X**^(2)^ are inputted to their respective autoencoders, converted feature matrices **Y**^(1)^ and **Y**^(2)^ (shorter notations for **Y**^(1,2)^ and **Y**^(2,1)^) are used to compute an Optimal Transport plan across the two modalities. The IOT loss ℒ_*IOT*_ computed thanks to the transport plan and the regularization loss ℒ_*reg*_ constituting together the rIOT constraint, are used to enforce the alignment of modalities in the shared latent space. (b) Example of output of scConfluence, cell embeddings visualized using 2D projections and clustered to discover new cell subpopulations; (c) Other example of output of scConfluence, the cell embeddings can be used to impute features across modalities.

scConfluence makes use of both the original data **X**^(*p*)^ and the converted data **Y**^(*p, p*′)^ to learn low-dimensional cell embeddings 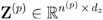 in a shared latent space of dimension *d*_*z*_. These embeddings can then be used for visualization and clustering, useful for discovering subpopulations of cells, and for imputation of features across modalities (Figure 1B-C).

For each modality *p*, scConfluence trains an autoencoder *AE*^(*p*)^ on **X**^(*p*)^ using modality-specific architectures^35^ and reconstruction losses 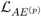 in order to retain all the complementary information brought by each modality. *AE*^(*p*)^ also performs batch correction by learning cell embeddings independent from their experimental batches of origin (see Methods). While frameworks based on autoencoders have been already designed in the context of diagonal integration^29,31,36^, the innovation of scConfluence is the combined use of Optimal Transport and regularized Inverse Optimal Transport (rIOT) for aligning cells in the shared latent space. Optimal transport (OT) is a mathematical toolkit for comparing high-dimensional point clouds^37^ that is gaining traction for addressing various problems in single-cell genomics: single-cell multi-omics cell matching^38,39^, paired multi-omics integration^20,39^, trajectory inference^39–42^ and predicting single-cell perturbation responses^43^. Solving the OT problem produces a correspondence map, i.e. transport plan, between point clouds based on their relative positions (see Methods). rIOT aims at addressing the inverse problem by inferring the relative positions of points based on a given transport plan^32^. scConfluence makes an innovative use of both OT and rIOT by first solving an OT problem leveraging weakly connected features (**Y**^(*p, p*′)^ and **Y**^(*p*′, *p*)^) to find a transport plan **P**^(*p, p*′)^ across modalities and then using rIOT on **P**^(*p, p*′)^ to adjust the cell embeddings inferred by *AE*^(*p*)^ and *AE*^(*p*′)^.

In more details, we first use **Y**^(*p, p*′)^ and **Y**^(*p*′, *p*)^ to compute a distance matrix between cells from different modalities which we then leverage to find an Optimal Transport plan 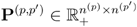 (see Methods). **P**^(*p, p*′)^ provides a correspondence map between cells of modalities *p* and *p*′ which we aim to leverage to determine the relative positions of cell embeddings in the shared latent space. This specific goal corresponds to the rIOT problem that we described above. In scConfluence, this is achieved by minimizing the loss 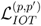 which penalizes distances between rows of **Z** ^(*p*)^ and **Z**^(*p*′)^ which are coupled by **P**^(*p, p*′)^. See Methods for a more formal explanation of the connection between our approach and rIOT. While 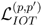 leverages biological prior knowledge to attract corresponding cells across modalities, it is not always sufficient to completely overlap them in the shared latent space. To address this, we add to the loss, as a regularization term, the unbalanced Sinkhorn divergence^44^ between the cell embeddings of each pair of modalities 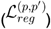. 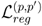, based on OT, is frequently used in machine learning to minimize the distance between high dimensional point clouds (see Methods). The gradients of both 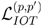 and 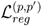 are back-propagated through the modality encoders in order to improve the across-modality alignment of cell embeddings. In addition, by using Unbalanced Optimal Transport in both 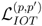 and 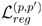, we do not force all cells to align, thus allowing scConfluence to deal with cell populations present only in one modality (see Methods).

The final loss optimized over the parameters of the *AE*^(*p*)^ with stochastic gradient descent is thus:

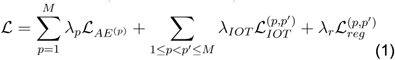

scConfluence separately uses all original features for dimensionality reduction in order to retain all the complementary information brought by each modality and leverages common information under the form of connected features to align cells with rIOT. Therefore, our innovative combined use of OT and rIOT allows scConfluence to avoid the loss of biological information generally resulting from modality conversion in state-of-the-art methods. As a consequence, scConfluence is much more robust to integration problems where very few features are connected across modalities (e.g. scRNA-surface protein data integration). In addition, the quality of scConfluence’s modality alignment depends on the transport plan ***P***^(*p, p*′)^ which relies only on the relative distances derived from the converted data **Y**^(*p, p*′)^ and **Y**^(*p*′, *p*)^. As a consequence, scConfluence can better deal with situations where strong batch effects between modalities are present in the converted data space. Furthermore, while state-of-the-art methods strictly enforce the complete mixing of cells across modalities, scConfluence, through the use of unbalanced OT, can cope with large discrepancies between the cell populations present in each modality. scConfluence is thus able to integrate single-cell modalities even when they do not contain the same cell types.

We extensively benchmarked scConfluence against five state-of-the-art methods: Seurat (v3.0), Liger, MultiMAP, Uniport and scGLUE^27–31^. Seurat, Liger and MultiMAPare widely used single-cell unpaired multi-omics integration methods in the computational biology community. Uniport is the main alternative to our method also using OT. Finally, scGLUE is the most recent and best performing method in the NeurIPS challenge on Open Problems in Single-Cell Analysis^45^.

### scConfluence outperforms the state-of-the-art on the integration of unbalanced cell populations

One of the main challenges of diagonal single-cell multi-omics integration is the need to deal with unbalanced cell populations. This requires aligning shared cell populations, independently of their size, and preserving modality-specific ones. We thus benchmarked scConfluence with the state-of-the-art based on its ability to integrate single-cell modalities sharing only a fraction of cell populations. As using simulated data based on distributional assumptions would favor methods making the same assumptions, we here designed a benchmark using scCATseq data profiled from HeLa, HCT and K562 cancer cell lines^15^. The choice of these data comes from the need to work with well-separated clusters, for which cell lines are an ideal example. In addition, having an equivalent proportion of cells per cluster in the two modalities allows us to design scenarios with different levels of unbalanceness in the cell populations. Of note, while scCATseq provides a joint profiling of scRNA and scATAC from exactly the same cell, the cell pairing information has not been used here as input of the various methods. To then test to which extent unbalanced cell populations affect the results of diagonal integration we modified the scCATseq data to represent three realistic situations: (i) removing half of K562 scRNA cells; (ii) removing all K562 scRNA cells and (iii) removing completely K562 scRNA cells and HCT scATAC cells. See Figure 2A for a schematic representation.

**Figure 2.**
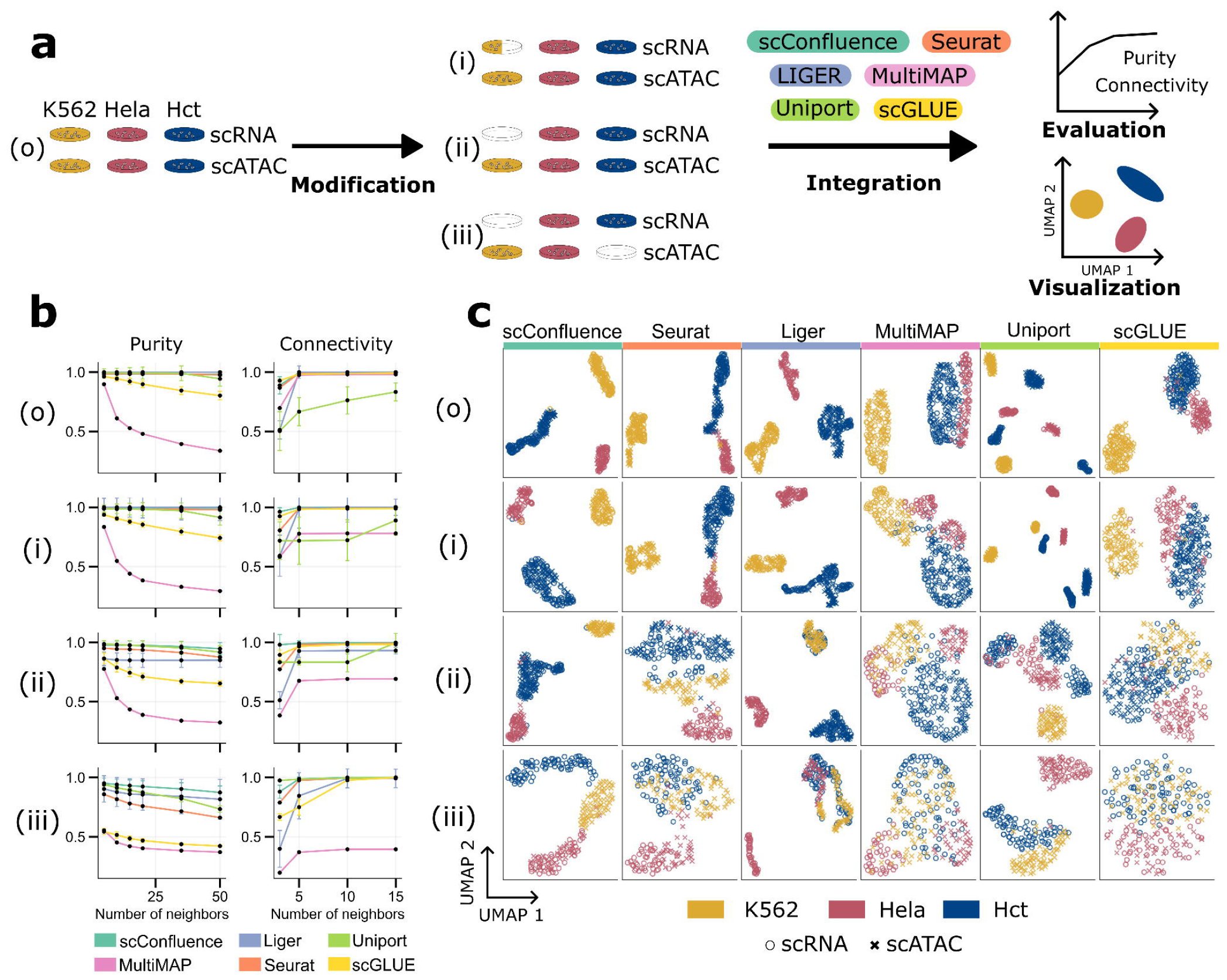
Benchmarking cell embeddings in unbalanced cell lines. (a) Schematic representation of the benchmarking process. Four scenarios are here considered: removing half of K562 scRNA cells, removing all K562 scRNA cells and removing completely K562 scRNA cells and HCT scATAC cells; (b) Purity and Connectivity scores are here reported for the six benchmarked methods (scConfluence, Seurat, Liger, MultiMAP, Uniport and scGLUE) on the four controlled settings derived from the cell lines data as described in panel a. Since purity and connectivity scores are based on nearest neighbors graphs, the plots report their behavior for various sizes of neighborhood (x-axis). Error bars in the plots specify the standard deviation across multiple random seeds for each method; (c) The six columns of this panel provide UMAP visualizations for the six benchmarked methods (scConfluence, Seurat, Liger, MultiMAP, Uniport and scGLUE) on the same four controlled settings derived from the cell lines data. Different colors in these UMAP plots correspond to the three different cell lines present in the data while the shape of the point markers correspond to the modality of origin of each cell (scRNA, scATAC).

We then benchmarked methods based on two main criteria: (i) their ability to group cells based on their cell line of origin (i.e. purity score^20^) and (ii) their capacity to mix modalities profiled from the same cell line (i.e. graph connectivity score^46^). See Methods for details.

As expected, all methods showed decreasing performances when the scenarios became less balanced. scConfluence outperformed the state-of-the-art in all scenarios, proving more robustness to variabilities in cell populations’ proportions (see Figure 2B-C). For the remaining methods, MultiMAP and scGLUE struggled the most to group cells based on their cell line of origin, while MultiMAP and Uniport were less performant in mixing modalities from shared populations. This can be observed also in the UMAP plots (Figure 2C). Regarding LIGER, the results here displayed concern its performances once setting the number of latent dimensions to three. This choice particularly advantages the method, whose performances get detrimental once a more standard value of latent dimensions is used (see Supp Figure 1). In addition, even when using three latent dimensions, LIGER displays higher variability in purity score across different runs, with respect to all other methods.

### scConfluence outperforms the state-of-the-art in scRNA-surface protein and scRNA-scATAC integration

To then benchmark scConfluence vs the state-of-the-art on larger and more realistic diagonal integration scenarios, we considered two 10X Genomics Multiome (scRNA+scATAC) datasets: (i) *PBMC 10X*, a human PBMC dataset with 9,378 cells per modality (ii) *OP Multiome*, a human bone marrow dataset, with 69,249 cells per modality profiled from different sites and donors constituting a total of 13 batches^47^; plus two CITE-seq (scRNA+surface protein) datasets: (i) *BMCITE*, a human bone marrow dataset with 30,672 cells per modality where 23 surface protein levels were measured^27^ (ii) *OP Cite*, a human bone marrow dataset with 90,261 cells per modality profiled from different sites and donors constituting a total of 12 batches and with 134 surface proteins^47^. These are gold standard datasets in multi-omics integration, already used to benchmark state-of-the-art methods^27,31,45,47^. We chose paired multi-omics data to test diagonal integration in order to have ground-truth matching between cells, useful for evaluating the performances of the various methods. Of note, the data have been treated as unpaired by the various methods and the cell pairing information has only been used for performance evaluation. In addition, the data are provided with high-quality cell labels useful for performance evaluation. For details on the data see Supplementary Table 1 and for their preprocessing see Methods.

A successful integration method should: (i) produce biologically meaningful integrated cell embeddings, i.e. organizing cells according to cell types and states, and (ii) align cells profiled from different modalities (e.g. scRNA, scATAC) that are paired or at least from the same cell type/state. We used purity score to evaluate (i), as done in the previous section. For (ii), we used two scores: Fraction Of Samples Closer Than the True Match (FOSCTTM), to evaluate the closeness of paired cells, and transfer accuracy^48^, to measure the proximity between corresponding cell types across modalities in the shared latent space (see Methods). Concerning MultiMAP, its output used for downstream analyses is a neighborhood cell graph only encoding closest interactions. This link thresholding in the neighborhood cell graph results in artificially low performances with FOSCTTM. For this reason, FOSCTTM was not reported for MultiMAP.

Regarding scRNA-scATAC integration (Figure 3B), scConfluence is the best performing method, leading in two out of three evaluation scores (Purity and Transfer accuracy). Concerning FOSCTTM, scGLUE has the best performances, immediately followed by scConfluence and Uniport. All methods perform better on *PBMC 10X* than *OP Multiome*. This is not surprising as *OP Multiome* contains more cell populations and strong batch effects, corresponding to several donors and sequencing sites. Of note, on this dataset, scConfluence performs best for batch correction (Supp Figure 2). Overall, for scRNA-scATAC integration, scConfluence is the method achieving the best compromise between producing a biologically meaningful integrated cell embedding and aligning cells profiled from scRNA and scATAC. In Figure 3C, UMAP visualizations illustrate the quality of the integration results obtained by scConfluence, with respect to the mixing of the modalities, the correction of batch effects and alignment of annotated cell types. For all other methods see Supp Figure 3-4.

**Figure 3.**
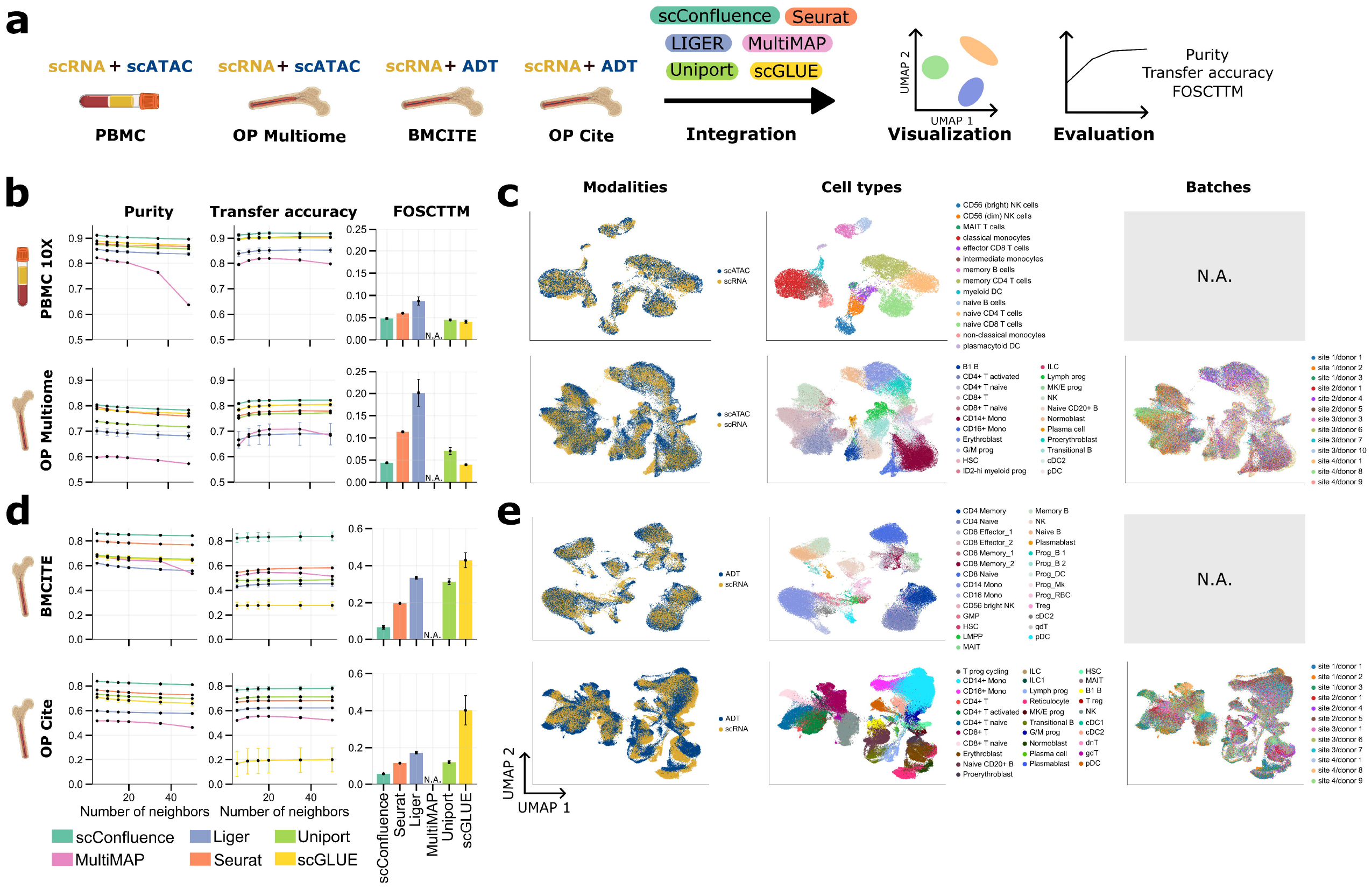
Cell embedding benchmark in gold-standard scRNA-surface protein and scRNA-scATAC datasets. (a) Schematic representation of the benchmarking process; (b) Purity, Transfer accuracy and Fraction Of Samples Closer Than the True Match (FOSCTTM) scores for the six benchmarked methods (scConfluence, Seurat, Liger, MultiMAP, Uniport and scGLUE) in two scRNA-scATAC datasets profiled from PBMC and bone marrow; (c) UMAP visualizations of scConfluence’s cell embeddings in the same datasets as panel b. Cells are colored based on their modality of origin, their cell type annotation or their batch of origin (when multiple batches are present in the data), respectively; (d) Same scores and methods as panel b, but computed on the two scRNA-surface protein datasets of the benchmark profiled from bone marrow; (e) UMAP visualizations of scConfluence’s cell embeddings on the two scRNA-surface protein datasets with cells colored according to the same rules as panel c.

In scRNA and surface protein integration (Figure 3D) scConfluence largely outperformed the state-of-the-art based on all three metrics on both datasets. On *BMCITE*, the relative improvement of scConfluence with respect to the second best is 9% in purity, 45% in transfer accuracy (corresponding to over 30% of the cells better classified by our method) and 66% in FOSCTTM. The performance gap is smaller on *OP Cite*, but still sizable with a relative improvement of 10% in purity, 10% in transfer accuracy (corresponding to over 5% of the cells better classified by our method) and 50% in FOSCTTM. The observed gap can be explained by the need of state-of-the-art methods for a large number of connections between the features of different modalities. This is not the case when integrating scRNA and surface protein data. For instance, in *BMCITE*, only 23 features are connected between the two modalities. As a consequence, most state-of-the-art methods have to subset the scRNA features to 23 protein-coding genes, thus discarding most of the information contained in the data. Moreover, scGLUE also struggles to align modalities since its prior feature graph contains thousands of nodes but only 23 edges.

The quality of our integration is highlighted by the UMAP visualizations in Figure 3E. While on *BMCITE* the modalities are completely mixed, on *OP Cite* a non-perfect mixing can be observed for few cell types/states (e.g. reticulocytes, erythroblasts and lymphoid progenitors). However, the integration of *OP Cite* data is a particularly challenging task, where a good tradeoff needs to be found between overlapping cells from different data modalities, correcting batch effects in each modality and defining a biologically meaningful integrated cell embedding (i.e. organizing cells according to cell types and states). Based on the evaluation in Figure 3D, scConfluence is the method achieving the best tradeoff. All other state-of-the-art methods suffer more in at least one of these objectives (Supp Figure 5-6). For instance, LIGER completely overlaps the two modalities, but provides integrated cell embeddings less biologically coherent than scConfluence.

### scConfluence robustly integrates scRNA and smFISH from mouse cortex, predicting genes with relevant spatial patterns

The phenotypic behavior of a cell, i.e. the cell state, results from the joint activity of the molecular regulation inside the cell and the influence of neighboring cells. Working with gene expression across space (e.g. in tissue context) is thus crucial to better characterize cell states. However, the possibility to jointly measure at single-cell and high-throughput resolution both spatial position and gene expression is still rare^49^. At the same time, other existing data have important limitations. On one hand, spatial high-plex imaging data (e.g. smFISH^50–52^, starMAP^53^) are limited by the possibility of only measuring a few genes (∼100-1000 genes)^54^. On the other hand, scRNA sequencing allows to sequence the full transcriptome but breaks tissues apart thus losing the spatial information^1^. Integrating these two types of data is thus the best opportunity we have to shed light on the role of spatial context in cell state definition.

With this aim, we applied scConfluence to integrate two gold standard datasets profiled from the mouse somatosensory cortex: (i) smFISH data of 33 selected marker genes measured in 4530 cells^55^; (ii) Smartseq2 data of ∼20k genes (including the 33 of the previous dataset) measured across 3005 cells^56^. As shown in Figure 4A, two outputs of scConfluence have been considered: (i) cell embeddings, whose quality is evaluated based on the same criteria used above (except for FOSCTTM since the data is unpaired) and (ii) imputations of the expression levels of unmeasured genes in the smFISH experiment. scConfluence’s results are here compared with the same state-of-the-art methods as before, with the only addition of GimVI^57^ which was specifically designed for scRNA and spatial high-plex imaging data.

**Figure 4.**
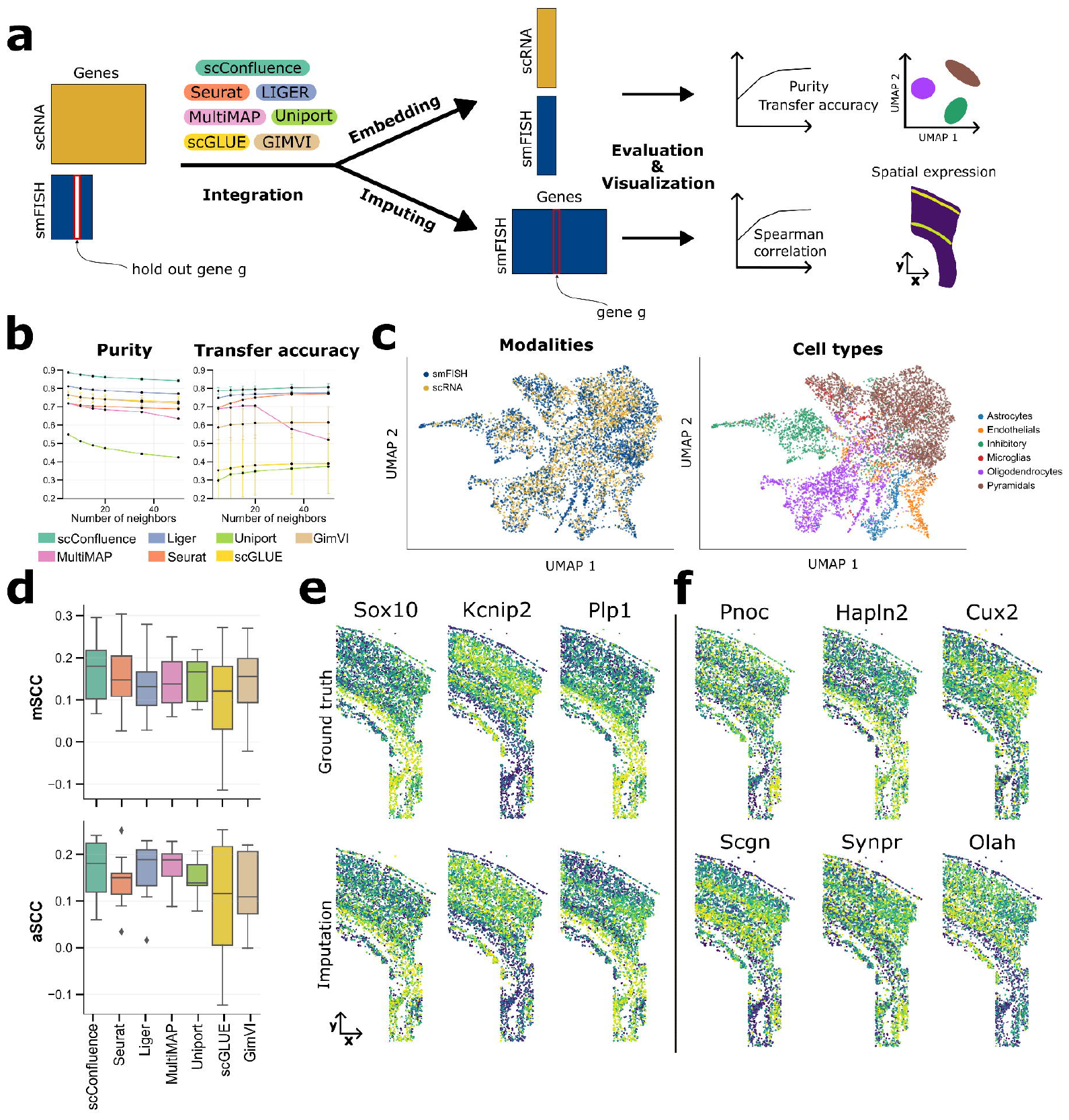
Cell embeddings and gene imputations resulting from scRNA and smFISH integration in mouse somatosensory cortex. (a) Schematic representation of the integration and imputation process; (b) Purity and Transfer accuracy scores of the seven benchmarked methods (scConfluence, Seurat, Liger, MultiMAP, Uniport and scGLUE, GimVI); (c) UMAP visualizations of scConfluence’s cell embeddings colored by the modalities of origin and their cell type annotations; (d) Boxplots of average and median Spearman correlation coefficients (aSCC and mSCC) (n = 11, no statistical method was used to predetermine sample size) between real and imputed smFISH genes. In the boxplots, the center line, box limits and whiskers denote the median, upper and lower quartiles and 1.5× interquartile range, respectively; (e) Spatial pattern of expression of scConfluence’s imputations (bottom) on three held-out smFISH genes and their ground-truth pattern of expression (top). (f) scConfluence’s imputed spatial pattern of expression of six scRNA genes not measured in the smFISH experiment.

Regarding the quality of cell embeddings, scConfluence outperforms all state-of-the-art methods according to both cell type purity and transfer accuracy (Figure 4B-C, Supp Figure 7). Thus, scConfluence proved again the ability to leverage a small number of common features to perform diagonal integration. Regarding the smFISH imputations, scConfluence enables us to predict features across modalities by connecting the smFISH encoder with the scRNA decoder. Indeed, the scRNA decoder can take as input a cell embedding from any modality and output its estimated scRNA profile. To evaluate the quality of the imputations, as done in^57^, we created multiple scenarios holding out ∼10% of the smFISH genes (see Methods). The proximity between the imputed and the ground-truth smFISH measurements was then calculated based on average and median Spearman correlations (aSCC and mSCC), as in^29^. The Spearman correlation is a natural choice for this task^29,57^ since it is less sensitive to outliers and focuses on the monotonic relationship (not necessarily linear) between pairs of observations. This is particularly relevant since we are interested in rewarding imputations which reflect the ground-truth’s pattern of expression rather than its absolute values. As shown in Figure 4D, gene imputation is very challenging, as aSCC and mSCC values are relatively low even for the most performant methods (median score around 0.1-0.2). Overall, according to mSCC, scConfluence outperforms the state-of-the-art methods, while according to aSCC scConfluence performs comparably to the best state-of-the-art methods. In Figure 4E, the quality of the imputations of scConfluence can be assessed also visually for the genes *Sox10, Kcnip2, Plp1* (all other genes are available in Supp Figure 8). The results suggest that scConfluence provides predictions spatially coherent with the ground-truth. In particular, *Sox10*, and *Plp1* exhibit higher expression in oligodendrocytes while *Kcnip2* displays higher expression in excitatory and inhibitory neurons (see^55^ for brain region annotation).

In addition, for the genes measured in scRNA but not in the smFISH data, scConfluence predicted some interesting spatial patterns (Figure 4F). In particular, for *Pnoc, Hapln2* and *Cux2*, known markers of inhibitory neurons, oligodendrocytes and upper neuronal layers respectively, scConfluence imputed smFISH profiles coherent with existing studies^56,58^. Finally, scConfluence also suggests additional genes having interesting spatial patterns: *Scgn*, highly expressed in the excitatory neurons from layers 4 and 6, *Synpr*, highly expressed in the region corresponding to the caudoputamen, and *Olah*, highly expressed in hippocampal and layer 6 neurons. These last results prove the ability of scConfluence to provide new relevant biological hypotheses to be followed-up experimentally.

### scConfluence integrates highly heterogeneous scRNA, scATAC and cyTOF leveraging their complementarity to improve cell type identification in PBMCs

A crucial challenge in biology is to take advantage of the complementarity between different data modalities to achieve a better understanding of cellular heterogeneity. While this is easier to achieve when the data are profiled from the same set of cells (e.g. 10X Multiome, CITE-seq), it becomes more challenging on unpaired data. Here, we bring this challenge to its extreme by performing diagonal integration of three PBMC single-cell omics data profiled from different cells, different donors and by different laboratories. The aim is to test to which extent scConfluence takes advantage of the complementarity between different data modalities despite the significant across-dataset variations.

We thus applied scConfluence to the diagonal integration of three human PBMC datasets extracted in highly heterogeneous settings: (i) Seq-Well-based scRNA-seq dataset of 16627 cells^59^; (ii) 10x Genomics scATAC-seq (Chromium platform) dataset of 21261 cells^60^ and (iii) single cell resolution mass cytometry (Helios CyTOF system) dataset where 48 proteins were measured in 43232 cells^61^. This configuration is particularly challenging for diagonal integration as in most real applications the different modalities would have been extracted from a single group of donors in comparable conditions, a situation characterized by much lower biological and technical variations.

For each of the three datasets, cell type annotations were provided in their original publication. Strong discrepancies could be observed in the depth of annotation of most of the cell types. For example, B cells in scATAC are divided into naive, memory and plasma; in CyTOF instead they are divided into naive, memory and double negative and in scRNA they are merged in a single B cell population. In addition, some cell types were modality-specific, for example MAIT T cells for CyTOF, plasma cells for scATAC data. Such discrepancies might be due to the absence of such cell types in some modalities, to their misclassification or to differences in annotation depth in the original studies.

scConfluence successfully integrated all three modalities in a common latent space where cells were organized according to cell types and states independently from their modality of origin (see Figure 5B-E). Indeed, as it can be already observed from the UMAP of the three omics integration (Figure 5B-D), cells from different modalities and corresponding to the same cell type annotation overlap in the latent space. In addition, once clustering cells in the integrated latent space (Figure 5F), the obtained clusters are consistent with the annotations of each modality (see Figure 5G-I). However, our integrative analysis also provides additional information (Figure 5F-I). The cells annotated as B cells in scRNA are split into three clusters from the three omics integration (Figure 5G, clusters: 0, 1, 2). In scATAC (Figure 5H) these three clusters correspond to cells annotated as memory, naive and plasma B cells. Similar conclusions can be derived from the CyTOF annotation (Figure 5I). We can thus assume that the cells classified in scRNA as cluster 0-2 also correspond respectively to memory, naive and plasma B cells. scConfluence’s integration thus had a crucial role in re-annotating the scRNA B cell cluster into appropriate subpopulations. We then further verified whether this subclustering of B cells in scRNA corresponds to real biological signal or to the random splitting of scRNA B cells driven by the artificial mixing of cells across modalities. With this aim, we identified the differentially expressed genes in clusters 0-2 for both scRNA and scATAC-derived gene activity, separately. CyTOF was excluded from this analysis because of the low number of features (only 48 proteins). We then tested the significance of their intersection (see Methods, Figure 5F, Supp Table 2), finding an overlap of: 30 genes (corresponding to a -log10FDR of 19) for cluster 0, 12 genes for cluster 1 (corresponding to a -log10FDR of 10) and 232 genes for cluster 2 (corresponding to a -log10FDR of 37). All of them being well beyond the standard FDR threshold of 0.01 proves that clusters 0-2 share the same differentially expressed genes in scRNA and scATAC. In addition, the common differentially expressed genes contain known markers of memory, naive and plasma B cells: *AIM2* and *RALGPS2*^22^ for memory B cells; *BTG1, TCL1A* and *YBX3*^22^ for naive B cells and *MCL1*^62^ for plasma B cells. Taken together these results thus confirm that the splitting of scRNA cells annotated as B cells into three subclusters (0-2), is not the result of an artificial modality alignment, but corresponds to real biological signals not identified in the previous unimodal scRNA analysis^59^.

**Figure 5.**
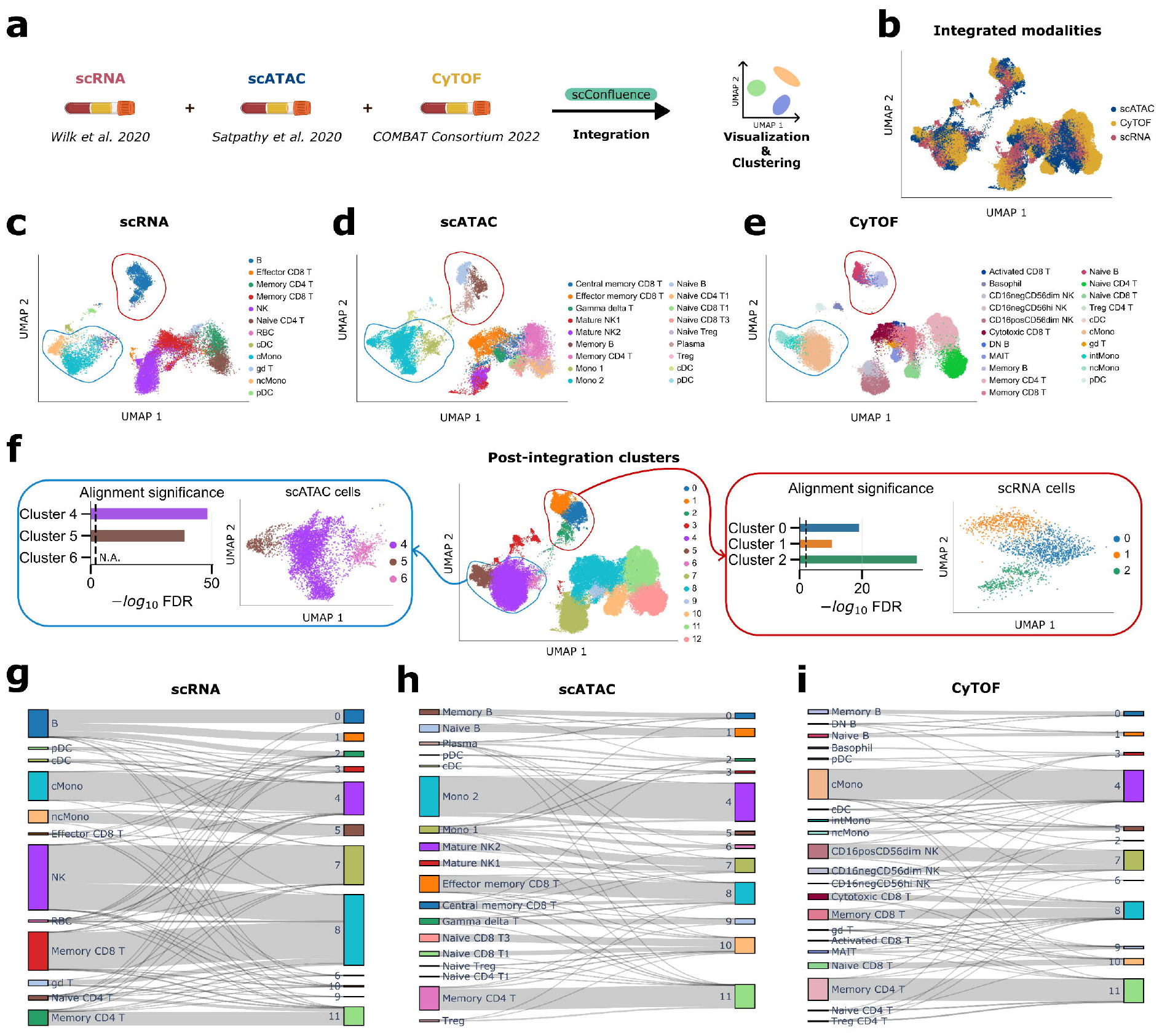
Tri-omics integration and sub-clustering of PBMC data. (a) Schematic representation of the integration; (b) UMAP visualization of all the integrated cell embeddings colored by their modality of origin; (c-e) UMAP visualization of scConfluence’s integrated cell embeddings plotted one modality at a time and colored by their cell type annotation of origin. The red circles highlight B cells which are already sub annotated in scATAC and CyTOF. The blue circles highlight monocytes which are already sub annotated in scRNA and CyTOF; (f) UMAP visualization of all the integrated cell embeddings colored based on inferred cluster annotations. Additional plots are provided for ATAC monocytes and RNA B cells which have been sub-clustered. The significance of the overlap between the marker genes obtained from scRNA and scATAC for each sub-cluster (Fisher’s exact test) is plotted. The dashed vertical line corresponds to FDR = 0.01. No alignment significance score is reported for cluster 6 as it only contains cells from the scATAC experiment; (g-i) Sankey diagrams displaying the comparison between cell annotations in their original publication and in our integrative analysis.

B cells are not the only example of cell populations benefitting from single-cell multi-omic integration. Monocytes are also annotated differently across single-cell omics data. Indeed, the scRNA study clusters them into classical and non-classical; CyTOF divides them into classical, non-classical and intermediate; scATAC splits them into Mono 1 and Mono 2. scConfluence’s integration of these three omics data divides monocytes into three clusters (4, 5 and 6), 4 and 5 having a good correspondence with classical and non-classical monocytes, respectively (see Figure 5G,I). As shown in Figure 5I, intermediate monocytes tend to cluster in the shared latent space together with non-classical monocytes (cluster 5), probably due to the fact that the clustering algorithm is splitting cell populations into discrete groups while this is a continuum of cells. In addition, the Mono 2 population of scATAC is split into clusters 4 and 5, thus containing both classical and non-classical monocytes. On the opposite, cluster 6 only corresponds to Mono 1 from scATAC, possibly representing a different state of monocytes not fitting within the classical/non-classical subdivision. To confirm such conclusions, we ran the same statistical test as earlier (Figure 5F, Supp Table 3) and found an intersection of differentially expressed genes between scRNA and scATAC of 226 genes for cluster 4 (corresponding to a -log10FDR of 48) and 80 genes for cluster 5 (corresponding to a -log10FDR of 39). In addition, the shared differentially expressed genes contained *CD14*, known marker of classical monocytes, for cluster 4 and *CD16*, known marker of non-classical monocytes, for cluster 5. Concerning cluster 6, composed only of scATAC cells, the overexpression of *CD2* and *CCR7* (log2 fold change of 5.61 and 5.40 respectively) could be observed, possibly suggesting that cluster 6 is a group of monocytes transitioning into Dendritic Cells^63,64^ (see Supp Table 4).

### scConfluence integrates scRNA and neuronal morphologies highlighting morphological heterogeneity in neuronal cell types of mouse motor cortex

The experiments above were focused on molecular data (e.g. transcriptomics, epigenomics and proteomics), but single-cell analysis can also benefit from other data modalities, such as imaging. A classical situation where imaging data play a key role is the study of neurons. Indeed, morphology imaging data provide a different classification of neocortical neurons with respect to scRNA data. An example of classification based on manual annotation of morphologies divides mouse neocortical interneurons into 15 groups^65^ representing different subgroups of Martinotti, neurogliaform, basket, single-bouquet, bitufted, bipolar, double-bouquet, chandelier cell, shrub, horizontally elongated, pyramidal and deep-projecting. On the other hand, in scRNA mouse motor cortex neurons have been classified into 90 populations^66^, corresponding to different subpopulations of Lamp5, Sncg, Vip, Sst, Pvalb pyramidal tract, near-projecting, Cortico Thalamic (CT), Extra Telencephalic (ET) and Intra Telencephalic neurons (IT). The integration of these two data modalities has thus a crucial role in unraveling neural heterogeneity and its associated biological functions^67^. This is an extremely challenging task that could not be tackled by the other state-of-the-art methods, as no natural connection exists between the pixels of an image and the features of scRNA data (i.e. genes).

We considered a dataset of 1214 adult mouse primary motor cortex cells profiled with Patch-seq, providing scRNA-seq, neuronal morphologies and electrophysiology measurements. The dataset is classified, based on scRNA, into Lamp5, Sncg, Vip, Sst, Pvalb, CT, ET and IT neurons extracted from layers 1, 2/3, 5 and 6^68^. Out of the 1214 cells, only 625 cells were profiled for both scRNA and morphologies, while for the remaining 589 cells only scRNA was available. This is not surprising as Patch-seq is difficult to master, thus implying the production of data containing some modalities and missing others, typical scenario of interest for diagonal integration. As shown in Supp Figure 9A, cells from scRNA perfectly organize according to the cell labels obtained in^68^. On the contrary, Supp Figure 9B shows that the scRNA labels do not fully capture the heterogeneity present in the morphology data, thus further suggesting that this modality contains complementary information. We thus investigated the role of such complementarity, by integrating with scConfluence the 625 available morphologies together with the 589 scRNA profiles (Figure 6A). The scRNA profiling of the first set of cells has been used to bridge the two modalities. This means that genes from scRNA have been considered as the connected features. These measurements are ideal to compute a reliable transport plan across the modalities as they come from the same sequencing technology and dataset.

**Fig 6.**
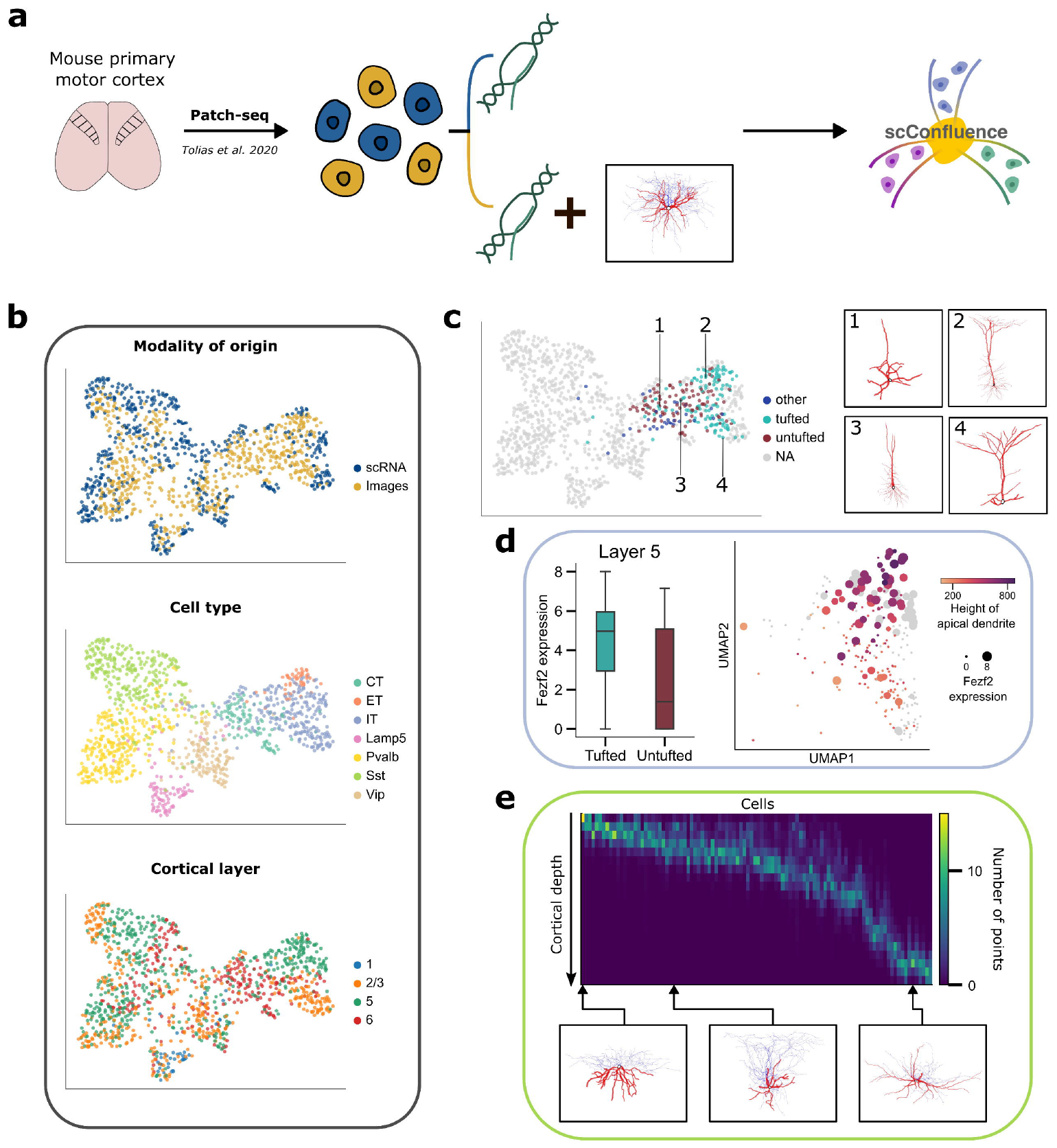
Integration of scRNA-seq and neuronal morphologies in the mouse primary motor cortex. (a) Schematic representation of the integration; (b) UMAP visualizations of the integrated cell embeddings colored by their modality of origin, their cell type annotations and their cortical layers of origin; (c) UMAP visualization of the integrated cell embeddings colored by their morphological labels which are only available for excitatory neurons. The terms ‘tufted’ and ‘untufted’ correspond to visual inspection of the neurons’ apical dendrites; some examples of neuronal morphologies are displayed next to the UMAP plot; (d) Pattern of expression of *Fezf2* in IT neurons. The boxplot on the left shows the distribution of expression of *Fezf2* in untufted and tufted IT neurons from layer 5. The UMAP plot of IT neurons shows the correlated pattern of variation of *Fezf2* expression (corresponding to the size of the points) and the height of apical dendrites (corresponding to the color gradient); (e) Heatmap representing the depth profiles of Sst neurons’ axons perpendicular to the pia. Cells have been sorted based on the depth of their soma.

The cells in scConfluence’s shared latent space were broadly organized according to the previously defined scRNA populations (Figure 6B). At the same time, morphological heterogeneity could be detected in some of these populations. For example, as shown in Figure 6C, excitatory neurons (CT, ET, IT) are organized into three morphological categories: “tufted”, “untufted” and “other” based on the visual inspection of their apical dendrites^69^. Most of the CT neurons are untufted and other, ET neurons are mainly tufted, finally, IT neurons result in a continuum progression from tufted to untufted. This progression seems associated with their layer of origin. For example, tufted IT neurons tend to be from layers 2/3 and 5, while untufted IT neurons are mostly from layer 6. Such morphological heterogeneity is extremely relevant as the geometry of tuft dendrites has an impact on the integrative properties of excitatory neurons^70–72^. In addition, we observe a higher expression of the Transcription Factor *Fezf2* in tufted IT neurons from layer 5 (see Figure 6D). This result is concordant with the hypothesis that *Fezf2* expression is required for the maintenance of tuftness in IT neurons^73,74^. However, we also observe tufted cells not expressing *Fezf2* as well as untufted cells expressing *Fezf2*, thus raising the possibility that other factors might be involved in such a process. Focusing then on all IT neurons, both the expression of *Fezf2* and the length of apical dendrites display a continuous gradient along the same one-dimensional manifold (Figure 6D). In agreement with this, both *Fezf2* activity and length of apical dendrites have been independently found to be highly correlated with calcium signaling^75,76^, which is connected to dendritic excitability through calcium electrogenesis^77,78^. Our observation has particular biological relevance as it could represent not only a simple association, but a causal effect of *Fezf2* on the morphology of IT neurons resulting in a regulation of dendritic excitability. This hypothesis is supported by the fact that *Fezf2* has been already shown to play a key role in the determination of the function, dendritic morphology and molecular differentiation of CT neurons^79^.

Furthermore, Somatostatin-expressing neurons (Sst), which are known to be morphologically diverse^80^, seem to be organized according to their layer of origin, with layer 2/3, layer 5 and layer 6 moving from left to right in the last UMAP plot of Figure 6B. This laminal organization is associated with a morphological pattern of variation, as shown by the axonal depth profiles in Figure 6E. In layer 2/3 we observe a higher presence of Martinotti cells extending their axons up to layer 1. Indeed, Martinotti cells are known to make contacts in layer 1 with the distal tuft dendrites of pyramidal cells^81^. On the other hand, deeper layers contain more non-Martinotti cells which seem to often target neurons inside their own layer.

## Discussion

The impressive abundance of unpaired multimodal single-cell data has motivated a growing body of research into the development of integration methods. However, the state-of-the-art suffers from two major drawbacks: (i) the loss of biological information due to across-modalities feature conversion and (ii) the presence of populations only profiled in one data modality.

We introduced scConfluence, a novel method for single-cell diagonal integration combining uncoupled autoencoders with regularized Inverse Optimal Transport (rIOT) on weakly connected features. scConfluence produces informative cell embeddings in a shared latent space by leveraging the complementarity of multiple modalities profiled from different groups of cells. This aim is achieved by using autoencoders on the full data matrices, allowing simultaneous dimensionality reduction and batch correction of different unpaired data modalities, together with rIOT on connected features to align cells in the shared latent space. This approach allows scConfluence to leverage prior knowledge without discarding the modality specific features which also provide relevant biological information.

Unlike the state-of-the-art, scConfluence does not rely on the assumption that most features are strongly connected across modalities. Indeed, as soon as such connections allow us to compute meaningful relative distances between cell populations the integration will be successful. This can be achieved even when there are few connected features, as in smFISH-scRNA integration, or when such connections are not perfect, as for proteins and scRNA integration^82^. In addition, the use of unbalanced Optimal Transport allows us to account for the presence of cell populations not shared across modalities.

We extensively benchmarked scConfluence in several scRNA-surface protein and scRNA-scATAC integration problems proving that it outperforms the state-of-the-art. We then explored scConfluence’s ability to tackle complex and crucial biological questions. First, we integrated with scConfluence scRNA and smFISH profiled from mouse somatosensory cortex and we imputed spatial patterns of expression for *Scgn, Synpr* and *Olah* relevant for future biological investigations. Second, scConfluence’s integration of scRNA-seq, scATAC-seq and CyTOF in highly heterogeneous human PBMC datasets refined the classification of B cells and Monocytes. Finally, through the integration of neuronal morphological images with scRNA-seq from the mouse primary motor cortex, scConfluence shed light on the combined impact of *Fezf2* expression and apical dendrite morphology on information processing in Intra Telencephalic neurons.

A challenging aspect for scConfluence and all the state-of-the-art is the need of properly dealing with rare cell populations. Indeed, rare populations are harder to detect as they are under-represented in parameter estimation. This is even more challenging for methods relying on mini-batch gradient descent (such as scConfluence, scGLUE and Uniport). Indeed, rare populations are much less likely to be simultaneously sampled from each modality in the mini-batches. At the same time, mini-batch optimization is necessary to scale to millions of cells. In addition, scConfluence, as much as all other state-of-the-art diagonal integration methods, relies on connections between features of different modalities. Such connections are not always available, as for example when integrating electrophysiology measurements with gene expression profiled from different neurons.

One of the main advantages of scConfluence is its modularity, allowing the users to choose their preferred unimodal dimensionality reduction method. For the modalities analyzed in this paper (scRNA-seq, scATAC-seq, CyTOF, smFISH, Patch-seq) ad-hoc autoencoders are proposed. However, for new modalities the users can choose whether to use a classical fully-connected autoencoder with the *L*_2_ loss or a more tailored solution available in the literature. Such a tailored solution could be a novel autoencoder architecture, or even any parametric dimension reduction model which can be optimized with stochastic gradient descent. Future developments could further improve the performances of scConfluence by plugging-in more advanced dimensionality reduction models recently developed or soon-to-be developed.

Regarding future perspectives, while this work is focused on unpaired multimodal data, paired multimodal data also start to increasingly accumulate. We can thus expect a relevant need for methods able to jointly integrate these two types of multimodal data. In this setting paired data would represent an extremely reliable prior knowledge to guide the alignment of unpaired cells. In addition, they could possibly bring new biological information, not already encoded in the single data modalities. Future developments of scConfluence should be aimed at tackling this intriguing emerging challenge.

## Methods

### Notations

For two vectors 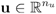 and 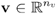, we use the notations:

(**u** ⊗ **v**)_*ij*_ = *u*_*i*_ + *v*_*j*_ and (**u** ⊕ **v**)_*ij*_ = *u*_*i*_ + *v*_*j*_. For two matrices **U** ∈ ℝ ^*n*×*d*^ and **V** ∈ ℝ ^*n*×*d*^ of identical dimensions, we’ll use the scalar product notation ⟨, ⟩ to denote the Frobenius inner product 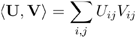.

### Optimal transport

Optimal Transport (OT), as defined by Monge^83^ and Kantorovich^84^, aims at comparing two probability distributions by computing the plan transporting one distribution unto the other with the minimal cost. While the OT theory has been developed in the general case of positive measures, our application only involves point clouds which are uniform discrete measures 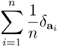 where the set of is the support of the point clouds. Therefore, to avoid adding unnecessary complexity in the notations we will denote the probability measures just as the set of positions **a**.

The classical OT distance, also known as the Wasserstein distance, between two point clouds 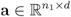 and 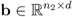 is defined as

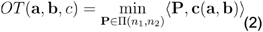

where 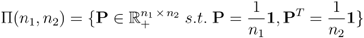 and *c* is a ground cost function used to compute the pairwise dissimilarity matrix 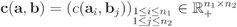 that encodes the cost of transporting mass from one point (e.g. cell) to another. In this uniform discrete case, the coupling **P** ∈ ∏ (*n*_1_,*n*_2_) is a matrix that represents how the mass in the point cloud **a** is moved from one point to another in order to transform **a** into **b**.

As real data often contains outliers to which OT is highly sensitive, a more robust extension of OT called unbalanced OT^33^ has been developed.

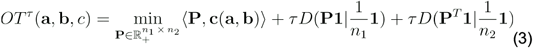

where *τ* is a positive parameter controlling the looseness of the relaxation.

In this formulation, the hard constraint on the marginals of the optimal plan is replaced with a soft penalization *D* which measures the discrepancy between the marginals of the transport plan *P* and the uniform distributions on **a** and **b**. While setting *τ* = + ∞ recovers the balanced OT problem (Eq. 2), using τ < + ∞ allows the transport plan to discard outliers and deal with unbalanced populations. Indeed, in (Eq. 3), unbalanced OT achieves a tradeoff between the constraint to conserve the mass by transporting all of **a** onto **b** and the aim to minimize the cost of transport. When an outlier is too costly to transport, it is therefore discarded from the plan. A classical choice for *D* is the Kullback-Leibler divergence. It is defined for two discrete probability distributions represented as vectors of probabilities **P** and **q** as 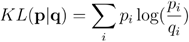. The Total Variation (TV) distance defined as *TV* (**p**,**q**) = ∣ *p*_*i*_ − *q*_*i*_ ∣ is also frequently used. The main difference between those two options is that when using TV, each point is either fully transported or discarded while using KL leads to transporting for each point a fraction of the mass which smoothly decreases as the cost of transport increases. We use both in different parts of our methods (see “Optimal Transport solvers”).

Adding an entropic regularization to the objective function of (Eq. 2) results in a new optimization problem noted as *OT*_*ε*_ (**a**,**b**,*c*), where *ε* is a positive parameter quantifying the strength of the regularization.

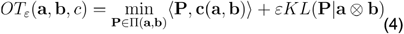

While setting *ε* = 0 recovers the unregularized OT problem (Eq. 2), using *ε* > 0 makes the problem - strongly convex. It can be solved computationally much faster than its unregularized counterpart with the GPU-enabled Sinkhorn algorithm^85^.

This entropic regularization can be used in the same fashion in (Eq. 3) to obtain the following problem:

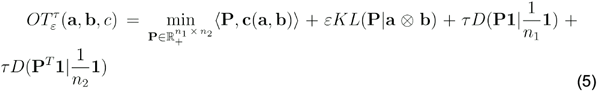

While 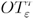 provides a scalable (thanks to the sinkhorn algorithm) and robust (thanks to the unbalanced relaxation) way to estimate the distance between point clouds, it shouldn’t be used as is for machine learning applications. Indeed, it suffers from a bias when *ε* > 0 and is not a proper metric for measures. In particular, 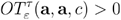. To solve this issue, a debiased version of (Eq. 5) has been introduced as the unbalanced Sinkhorn divergence^44^:

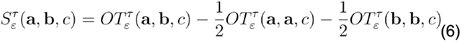

The Sinkhorn divergence 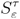 on the other hand is very well suited to define geometric loss functions for fitting parametric models in machine learning applications. Not only is it robust and scalable but it also verifies crucial theoretical properties such as being positive, definite, convex and metrizing the convergence in law.

To designate optimal transport problems, we’ll use the unified notations 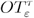 and 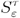 for all cases with τ = + ∞ referring to the balanced case and *ε* = 0 referring to the unregularized case.

### scConfluence

scConfluence takes as inputs data from *M* modalities with *M* ≥ 2 where each modality’s data comes under the form of a matrix 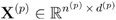 where the *n*^(*p*)^ rows correspond to cells and the *d*^(*p*)^ columns are the features that are measured in the p^th^ modality (e.g. genes, chromatin peaks, proteins). For each modality the vector **s**^(*p*)^whose entries are the batch indexes of the cells in **X**^(*p*)^ is also available. Additionally, for all pairs of modalities (*p,p*′) we have access to 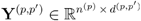 and 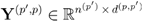 which correspond to **X**^(*p*)^ and **X**^(*p*′)^ translated to a common feature space. The method to obtain **Y**^(*p,p*′)^ for each modality is detailed later in the subsection “Building the common features matrix”.

ScConfluence leverages all these inputs simultaneously but in different components to learn low dimensional cell embeddings 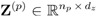 in a shared latent space of dimension *d*_*z*_. For each modality *p*, we use one autoencoder (AE) *AE*^(*p*)^ on **X**^(*p*)^ with modality-specific architectures and reconstruction losses 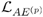, see the subsection “Training details”.

While variational autoencoders have become extremely popular in single cell representation learning, we decided not to use them. Indeed, variational autoencoders are trained by optimizing the ELBO which contains two terms, one for the reconstruction of the data and one which is the Kullback-Leibler divergence between the variational posterior and the prior distribution. This second term has been found to aim at a goal conflicting with the reconstruction and to lead to worst inference abilities^86^. With this in mind, we used classical autoencoders with an additional regularization. In our architecture, the encoder still outputs parameters of a gaussian with diagonal covariance as a variational model would, but instead of forcing this distribution to be close to an uninformative gaussian prior, we simply add a constant (0.0001) to the outputted standard deviation of the posterior distribution so that our model does not converge to a deterministic encoder during training. This stochasticity in the encoder acts as a regularization against overfitting as it forces the decoder to learn a mapping which is robust to small deviations around latent embeddings.

To handle batch effects within modalities, the batch information **s**^(*p*)^ is used as a covariate of the decoder as done in existing autoencoder-based methods for omics data^35^. Conditioning the decoding of the latent code z on its batch index s allows our AEs to decouple the biological signal from the sample-level nuisance factors captured in different batches.

Meanwhile, the **Y**^(*p,p*′)^ matrices are leveraged to align cells across modalities using Optimal Transport. For each pair of modalities (*p,p*′), we use the Pearson similarity (see Implementation details) to compute the cost matrix **c**_*corr*_ (**Y** ^(*p,p*′)^, **Y** ^(*p,p*′)^). Indeed, while the squared *L*_2_ distance is classically used in OT, the Pearson similarity has been shown to better reflect differences between genomic measurements^87^. Using this cost matrix, we derive the unbalanced Optimal Transport Plan **P** ^(*p,p*′)^ which reaches the optimum in 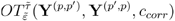. **P**^(*p,p*′)^ thus provides a partial plan to match corresponding cells from different modalities in the latent space. Using the unbalanced relaxation of OT to compute **P**^(*p,p*′)^ enables scConfluence to efficiently deal with cell populations present only in one modality. Indeed, cell populations which are not shared across modalities will have a higher transport cost and are more likely to be part of the mass discarded by the unbalanced OT plan. Once **P**^(*p,p*′)^ is obtained, it provides a correspondence map between modalities which determines which embeddings should be brought closer in the latent space. Since diagonal integration’s goal is to embed closely cells which are biologically similar, we enforce a loss term whose specific goal is this:

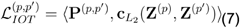

where 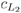 is the squared *L*_2_ distance such that 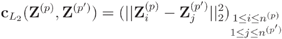.

Minimizing 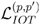 leads to reducing the distance only between the cell embeddings which are matched by **P**^(*p,p*′)^. We add to this loss a regularization term which reduces the global distance between the set of embeddings in **Z**^(*p*)^ and those in **Z**^(*p*′)^. This allows us to make sure that we do not only juxtapose corresponding cell populations from different modalities, but that they overlap in the shared latent space. To enforce this regularization, we use the unbalanced Sinkhorn divergence (Eq. 6) as both its computational and theoretical properties make it an ideal regularization function for our goal.

All those different objectives contribute together to the following final loss which we optimize over the parameters of the neural networks *AE*^(*p*)^ with stochastic gradient descent:

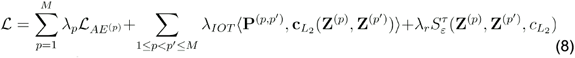

Where the *λp, λ*_*IOT*_ and *λ*_*r*_ are positive weights controlling the contribution of each different loss terms.

### Connection to regularized Inverse Optimal Transport

Our final loss (Eq. 8) can be decomposed in two main objectives, on one side the reconstruction losses whose goal is to extract the maximum amount of information out of each modality, on the other side the alignment loss 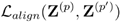, whose goal is to align cells across modalities in the shared latent space.

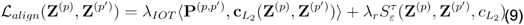

There is an intimate connection between 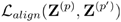 and the theory of Inverse Optimal Transport (IOT).

Regularized Inverse Optimal Transport (rIOT)^32^ refers to the problem of learning a pairwise dissimilarity matrix from a given transport plan **P** ∈ ∏ (*n*_1_, *n*_2_), with a certain regularization on C. In our case, it can be formalized as the following convex optimization problem:

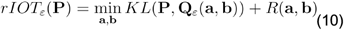

where **Q**_*ε*_ (**a**,**b**) is the balanced optimal transport plan achieving the optimum in 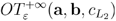 and *R* is a user-defined regularization. In our case, we want this regularization to force points coupled by **P** to completely overlap.

We prove that in the particular case of balanced plans, which corresponds to setting 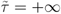 and *τ* = +∞ in our method, and with the regularizing function 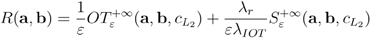, minimizing ℒ_*align*_ with respect to **Z**^(*p*)^ and **Z**^(*p*′)^ is equivalent to solving 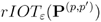. More formally, we prove that

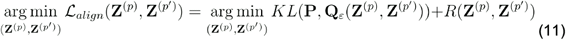

The proof uses the following lemma (See Supplementary Note 1).

#### Lemma

Let **a** and **b** be two point clouds of size *n*_1_ and *n*_2_ respectively. Given **P** ∈ ∏ (*n*_1_,*n*_2_) and denoting as **Q**_*ε*_ (**a**,**b**) the balanced entropic optimal transport plan achieving the optimum in 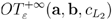, the following equality holds:

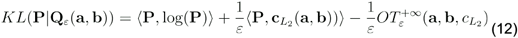

Using the lemma (Eq. 12) and the definition of *R* we prove (Eq. 11) by rewriting 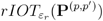 as:

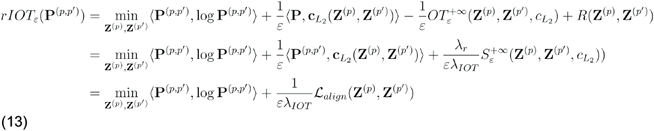

By noticing in (Eq. 13) that neither ⟨ **P**^(*p,p*′)^, log **P**^(*p,p*′)^ ⟩ nor the scaling factor 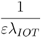 depends on (*Z*^(*p*)^, *Z*^(*p*′)^), we obtain (Eq. 11).

### Training details

#### Neural network architectures

The encoders and decoders are three-layer neural networks with ReLU activation function inspired by the architecture of the scVI VAE. We used a latent dimension of 16 for all datasets but adapted the number of neurons in hidden layers to the dimensionality of the datasets (see Supp Table 4). On scATAC and scRNA datasets which contained thousands of features, we did a first dimension reduction with PCA and used the 100 principal components as inputs of the encoder while the decoder outputted a reconstruction in the original feature spaces which was compared with the data prior to the PCA projection. For proteomic and smFISH modalities which contained much fewer features, we reduced the number of layers of both encoders and decoders to two. We used the same decoder architecture as scVI with the Zero Inflated Negative Binomial (ZINB) likelihood for the reconstruction loss on scRNA data. For other modalities however, we replaced the scVI decoder with a simple fully connected multi-layer perceptron and used the squared *L*_2_ distance as the reconstruction loss.

#### Optimal transport solvers

We used the Python package *POT* to compute the plans *P*^(*p,p*′)^ with the function ot.partial.partial_wasserstein. This implementation of unbalanced optimal transport uses the Total variation distance for the penalization of marginals. It is parameterized by the lagrangian multiplier *m* associated with 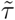 to control the unbalancedness of the plan. *m* is a parameter between 0 and 1 which quantifies how much mass is transported by the optimal plan. The use of TV to penalize the unbalanced relaxation allows **P**^(*p,p*′)^ to completely ignore cell populations which are identified to have no equivalent in other modalities. We set *m* = 0.5 and no entropic regularization 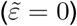 as POT’s CPU implementation was already fast enough on our mini-batches for us to afford avoiding using an approximation.

For the unbalanced sinkhorn divergence we used the python package *Geomloss*^88^ which has extremely efficient GPU implementations with a linear memory footprint. Indeed, while it cannot take as input a custom cost matrix as POT does, when the cost function is the squared *L*_2_ distance (as is the case for our regularization term) *Geomloss* uses KeOps^89^ to implement efficient operations with a small memory footprint and automatic differentiation. Geomloss uses the KL to penalize the unbalanced relaxation. We used the following hyperparameters: “p”= 2, “blur”= 0.01 (which corresponds to *ε* = 0.0001), “scaling”= 0.8, “reach”= 0.3 (which corresponds to *τ* = 0.09).

#### Training hyper parameters

All models were optimized using the Pytorch lightning library. We used the ADAMW optimizer^90^ with a learning rate of 0.003. The batch size was set to 256 times the number of modalities. 20% of the dataset was held out for validation and an early stopping was triggered when the validation loss didn’t improve for 40 epochs. 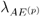 was set to 1.0 for all modalities except for ATAC where it was set to 5.0 due to the larger amount of content measured in the ATAC modality. The default value for *λ*_*IOT*_ was set to 0.01 while *λ* was set to 0.1 for *M* = 2 and to 0.03 for *M* = 3.

### Data preprocessing

#### scRNA preprocessing

We performed quality control filtering of cells on the proportion of mitochondrial gene expression, the number of expressed genes, and the total number of counts (using Muon’s filter_obs). Quality control filtering of genes was performed on the number of cells expressing the gene (using Muon’s filter_var). We then kept a copy of the raw counts data before applying the log-normalization which consists of normalizing counts for each cell so that they sum to 10000 (using Scanpy’s normalize_total) and then log transforming them (using Scanpy’s log1p). To subselect genes we took the union between the set of 3000 most variable genes in the normalized counts (using Scanpy’s highly_variable_genes with flavor=‘seurat’) and the set of 3000 most variable genes in raw counts (using Scanpy’s highly_variable_genes with flavor=‘seurat_v3’). Finally the log-normalized counts were used to compute the first 100 principal components which served as the input of the decoder while we kept a copy of the raw counts to evaluate the output of the decoder using the ZINB likelihood (except for the Patch-seq dataset where we used a fully connected decoder with the squared *L*_2_ loss on the log normalized counts).

#### scATAC preprocessing

We performed quality control filtering of cells on the number of open peaks and the total number of counts (using Muon’s filter_obs). Quality control filtering of peaks was performed on the number of cells where the peak is open (using Muon’s filter_var). We didn’t apply any further subselection of the peaks after the quality control. Cells were normalized using the TF-IDF normalization (using Muon’s tfidf). Finally the first 100 principal components of the normalized data were used as input to the encoder while the unreduced TF-IDF normalized data was used to evaluate the output of the decoder with a squared *L*_2_ loss.

#### Protein preprocessing (in Cite-seq and CyTOF)

Since the number of measured proteins is small and this data is less noisy than scRNA or scATAC, no quality control or feature selection was performed. We normalized the data using Muon’s implementation of the Center Log Ratio technique. This processed data was used for both the encoder and the decoder (with a squared *L*_2_ loss).

#### smFISH preprocessing

We performed quality control filtering of cells on the proportion of mitochondrial gene expression, the number of expressed genes, and the total number of counts (using Muon’s filter_obs). Quality control filtering of genes was performed on the number of cells expressing the gene (using Muon’s filter_var). For the smFISH gene counts we used the same normalization technique as in the original study: we normalized by both total number of molecules of all genes in each cell and the sum of each gene over all cells. This processed data was used for both the encoder and the decoder (with a squared *L*_2_ loss).

#### Patch-seq morphologies preprocessing

We retrieved the neuronal morphologies as 3D point clouds stored in .SWC files and did not have to do any quality control since only high quality morphologies could be reconstructed. We then used the *NeuroM* package^91^ to load the morphologies and project them onto the xy-plane (which is actually the xz plane since y and z were switched in the raw files) while coloring each point according to its neuronal compartment type (dendrites in red, axons in blue and soma in black). We then input those images in Google’s Inception v3 pre-trained deep neural network to extract features by retrieving the output of the last layer (with 2,048 dimensions). We then concatenated all these feature vectors in a matrix. This processed data was used for both the encoder and the decoder (with a squared *L*_2_ loss).

#### Building the common features matrix

The first step to construct the cross-modality cost matrices consisted in obtaining the **Y**^(*p,p*′)^ and **Y**^(*p,p*′)^ matrices.

- With scRNA and scATAC data this consisted in obtaining the gene activity matrix and subsetting the two matrices to the set of common genes. We obtained the gene activities using different techniques depending on the metadata available for each dataset. For the cell lines data we used Maestro^26^, for the Multiome PBMC data we used Signac^25^, the gene activities for the Open problems Multiome dataset had been already computed by the authors with Signac and for the tri-omics PBMC dataset we ran the R script provided by the authors on the github repository of their study https://github.com/GreenleafLab/10x-scATAC-2019/blob/master/code/04_Run_Cicero_v2.R using Cicero^92^.
- With scRNA and Protein data this consisted in manually inspecting the genecards website to find for each protein its associated coding gene and then subsetting the RNA and Protein data to the pairs available in both modality’s features.
- With scATAC and Protein we did the same as with RNA and Protein after obtaining the gene activities from ATAC.
- With RNA and smFISH since all genes measured in the smFISH experiment were also measured in the scRNA dataset we simply subset the scRNA genes to keep only the common genes.
- With RNA and Patch-seq morphologies since for both groups of cells we had access to the scRNA measurements we could directly use those as common features.

#### Building the biological cost matrix

Having obtained the converted data matrices **Y**^(*p,p*′)^and **Y**^(*p*′,*p*)^, we then applied to each modality’s data (ATAC gene activities were treated as RNA) the same preprocessing as described earlier. We then scaled both **Y** matrices (except for the Patch-seq since we were comparing scRNA data from the same dataset) and computed the cost matrix by using the correlation distance between each pair of cells from the two modalities using scipy’s *cdist*.

### Baselines

#### Seurat

We compare scConfluence to Seurat v3 as the v3 refers to the version aimed at tackling diagonal integration. In practice we used the R package *Seurat v4*.*3*.*0* which finds anchor pairs between cells from different modalities by searching for Mutual Nearest Neighbors after having reduced the dimension of the data with Canonical Correlation Analysis (CCA). Before running the CCA, all modalities are converted to the same features so we followed the same protocol as described above in the subsection “Building the common features matrix”, as it coincides with the indications described in the tutorials available in the Seurat documentation. We ran the Seurat method with default parameters, except for the Protein and smFISH datasets where we set the latent dimension to 15 since the default number of latent dimensions was close or even higher than the number of features measured. For gene imputation in the scRNA-smFISH experiment we used the TransferData function as indicated in the documentation.

#### LIGER

We compare scConfluence to Liger using the R package *rliger v1*.*0*.*0*. Liger relies on integrative non-negative matrix factorization (NMF) to perform diagonal integration and also requires as a first step to convert all modalities to common features. We did this step in the same way as for Seurat. For all datasets except the cell lines we ran Liger with default parameters. On the cell lines simulated experiment, using the default setting of 30 latent dimensions resulted in the embeddings from different modalities being completely separated. Since the latent dimensions can be interpreted as clusters in NMF we used this to set the number of latent dimensions to 3 which greatly improved Liger’s results. We could not tune other baselines similarly for this experiment as the dimension of their latent space can’t be interpreted similarly and this did provide a competitive advantage to liger since we used the knowledge that there were 3 main clusters in the dataset (which usually can’t be known when integrating new datasets). For the Protein and smFISH datasets we set the latent dimension to 15 since the default number of latent dimensions was close or even higher than the number of features measured. For gene imputation in the scRNA-smFISH experiment we used a knn regression with the scRNA embeddings serving as reference to predict the expression levels of held out genes for smFISH embeddings.

#### MultiMAP

We compare scConfluence to MultiMAP using the python package *MultiMAP v0*.*0*.*1*. MultiMAP is a generalization of the popular UMAP method^93^ to the unpaired multimodal setting. MultiMAP combines intra modality distances with prior knowledge-based cross modality distances to recover geodesic distances between all cells on a single latent manifold which can then be projected on ℝ^2^ for visualization. Intra modality distances are computed based on low dimensional projections of the data after preprocessing. We followed the documentation for this step although dimension reduction was not necessary for smFISH and proteomic datasets where the number of features was already lower than a hundred. To compute distances across modalities we converted pairs of modalities to a common feature space we did as described above in the subsection “Building the common features matrix”, as it coincides with the indications described in the tutorials available in the MultiMAP documentation. MultiMAP was run with default parameters on all datasets. For gene imputation in the scRNA-smFISH experiment we used knn regression with the scRNA embeddings serving as reference to predict the expression levels of held out genes for smFISH embeddings.

#### Uniport

We compare scConfluence to Uniport using the python package *Uniport v1*.*2*.*2*. Uniport uses one encoder which takes as input cells from all modalities converted to common features while using modality specific decoders to reconstruct each modality’s features. It also leverages unbalanced Optimal Transport in the latent space to force different modalities to mix in the latent space. For feature conversion we proceed as described above in the subsection *Building the common features matrix*, as it coincides with the indications described in the tutorials available in the Uniport documentation. We ran Uniport with default parameters on all datasets. For gene imputation in the scRNA-smFISH experiment we used the scRNA decoder to map the embeddings of smFISH cells to the scRNA domain as described in the Uniport documentation.

#### scGLUE

We compare scConfluence to scGLUE using the python package *scglue v0*.*3*.*2*. scGLUE simultaneously trains one variational autoencoder per modality and one graph variational autoencoder which learns feature embeddings based on a prior knowledge-based guidance graph containing connections between features from different modalities. We followed scGLUE’s documentation to construct the guidance graph for scRNA and scATAC integration. For scRNA and Protein integration where no documentation was available we created a graph where each coding gene was linked to its associated protein. For scRNA and smFISH integration we created a graph with links between each smFISH measured gene and the same gene in the scRNA data. We ran scGLUE with default parameters on all datasets, except for the Protein and smFISH datasets where we set the latent dimension to 15 since the default number of latent dimensions was close or even higher than the number of features measured. For gene imputation in the scRNA-smFISH experiment we used the scRNA decoder to map the embeddings of smFISH cells to the scRNA domain as in other autoencoder-based methods.

#### GimVI

We compare scConfluence to GimVI using the python package *scvi-tools v0*.*16*.*4*. GimVI is only applicable to scRNA and smFISH integration and simultaneously trains one autoencoder per modality while enforcing mixing between modalities in the latent space with a discriminative neural network trained in an adversarial way. We ran GimVI with default parameters and performed gene imputation as described in its documentation: we used the scRNA decoder to map the embeddings of smFISH cells to the scRNA domain.

### Evaluation metrics

We used several scoring functions to assess the quality of the embeddings provided by each method throughout the benchmarking. All methods were run with five different random seeds and we reported the median score, except for Seurat which contains no randomness and could therefore be run with one seed only. Apart from FOSCTTM, all metrics are based on the k-nearest neighbor graph of embeddings. To give a complete overview of the performance of the methods, we computed those metrics with *k* taking all values in {5,10,15,20,35,50}. Those metrics are therefore displayed as curves whose x-axis correspond to the values of *k*.

For the MultiMAP method whose output is not an embedding but a graph whose edge weights represent similarities between integrated cells, we can use this graph to compute nearest neighbors. Additionally, for the *OP Multiome* and *OP Cite* datasets which contained more than 60,000 cells per modality, we evaluated the methods after the training using a subset of 20,000 cell embeddings as the metrics were too expensive to compute on the full results of each method. We suppose in the following that there are only two modalities being integrated, as is the case for all benchmarked datasets.

#### Notations

For each cell *i* from the pth modality, we denote as 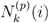 the *k* nearest neighbors of the cell’s embedding 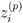 in the integrated latent space. *c*_*i*_ denotes the cell type label of cell *i*.

#### Purity

The purity score measures the average proportion of an integrated cell’s k-nearest neighbors that share the sample’s cell type annotation^20^. It thus varies between 0 and 1 with a higher score indicating a stronger performance.

The score can be written as 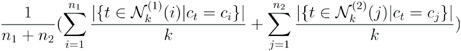.

#### Transfer accuracy

The transfer accuracy is the accuracy of a k-nearest neighbor classifier using one modality as reference and the other modality as query. Since both modalities can be the reference and the query, we compute the results of both classifications and report the average of the two scores.

#### Graph connectivity

The graph connectivity metric assesses how well cells with the same cell type label are connected in the kNN graph representation of the embeddings^46^. This score can be used to detect whether there exist discrepancies in the integrated latent space between cells from different modalities or experimental batches. For each different cell type *c*, we denote as *G*_*k*_(*c*) the subset of the integrated kNN graph containing only cells with label *c*. We compute for each cell type *c* the score *s*_*c*_ equal to the size of the largest connected component in *G*_*k*_(*c*) divided by the number of cells with label *c*. The final graph connectivity score is the average of the cell type scores *s*_*c*_.

#### FOSCTTM

The Fraction Of Samples Closer Than the True Match (FOSCTTM) metric has been used before to evaluate diagonal integration methods on paired multimodal datasets where both modalities are measured in the same cells^31,38,94^. Since this metric is only designed for paired datasets we can suppose that there are exactly cells for each modality and that they are ordered such that the ith cell in the first modality is the true match of the ith cell in the second modality. FOSCTTM aims at comparing for every cell from modality *p* the distance to its true match and the distance to all other cells in the opposite modality which we denote as *p*′ (since *p* ∈ {1,2}, *p*′ 3 − *p*). It is classically defined as:

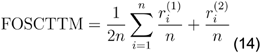

where 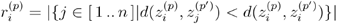

However, in this paper we compute it in a slightly different way. In the previous formula (Eq. 14), we replace 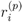 with 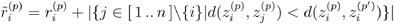. This means that rather than only assessing how close the true match of a cell’s embedding is compared to cells from the opposite modality, we assess how close the true match of the cell’s embedding is compared to all cells from both modalities.

With this formula we can simultaneously evaluate whether the mixing of the two modalities in the shared latent space is complete and verify that corresponding populations are accurately matched across modalities. Other metrics originally designed to assess batch effect correction are often used to evaluate the mixing of modalities such as the batch entropy of mixing^95^ but these don’t penalize artificial alignments. The complete overlapping of cells from different modalities only matters if those cells are biologically equivalent and this is assessed by our modified formulation of the FOSCTTM.

### Gene imputation

Both the scRNA-seq and smFISH datasets were downloaded using the scvi-tools helper functions *load_cortex()* and *load_smfish()* which select only the overlapping cell types to ensure the consistency of the imputation task. We then divided the 33 genes measured in the smFISH experiment into eleven disjoint groups of 3 genes. Each of this group corresponded to a different scenario where the three genes were removed from the smFISH data and held out and then imputed by each method.

Considering the prior knowledge using to connect features across modalities was extremely reliable (as we could map genes in the smFISH experiment with themselves in the scRNA-seq without any errors) we increased the weight of the IOT loss *λ*_*IOT*_ from 0.01 to 0.05 in this experiment.

For both methods which were autoencoder based, Uniport and scGLUE, we used the same technique as us to perform the imputation. For all other methods, we used knn regression using the scRNA-seq embeddings as reference to predict the expression levels of held out genes in smFISH embeddings.

We used the spearman correlation to quantify the similarity between the imputed values of a gene across all cells with the ground truth held out values. It is defined as the Pearson correlation between the rank values of those two vectors. As in the benchmarking section, we ran each method on each scenario with five initialization seeds (except Seurat which contains no stochasticity). For each gene in each scenario we kept the median spearman correlation across the five seeds. We then reported one score per imputation scenario and plotted the eleven scores as a violin plot. We can aggregate the spearman correlation of the three genes forming each scenario using the average or the median therefore we report both the average and median Spearman correlations (aSCC and mSCC).

For the visualization of the imputations, we made use of the recorded 2D positions of the smFISH cells to plot the cells as they are located in the tissue. To better visualize spatial patterns of the imputations, we used the histogram equalization technique on the imputed values.

### Tri-omics integration

We removed very rare cell types (containing less than 0.5% of the whole dataset) from all three datasets. this resulted in the removal of ATAC “Immature NK”, “Basophil”, “Naive CD8 T2” and “Naive CD4 T2”cells as well as CyTOF “Plasmablasts”, “cDC1”, “CLA+HLADR+ NK cells”, “Activated gd T cells”, “Unclassified”, “HLADR+CD38-CD4 T cells”, “Cytotoxic CD4 T cells”, “DN T cells” and “Activated CD4 T cells”.

We clustered the cell embeddings from all modalities using scanpy’s louvain with a resolution of 0.5. We then focused on the B cells and monocytes clusters which we reclustered with a resolution of 0.2.

To assess whether the subclusters we found were correctly aligned across modalities we used the same methods as described in scGLUE^31^ except that we only used scRNA and scATAC derived gene activities. Indeed, including the CyTOF data in this analysis would have resulted in removing too many features to be able to design a statistical test with sufficient power. For each of the two populations we subclustered (B cells and monocytes), we tested for significant overlap in cell type marker genes. For both gene expression and gene activities, the cell type markers were identified using scanpy’s one-versus-rest Wilcoxon rank-sum test with the following criteria: FDR < 0.05 and log fold change > 0. The significance of marker overlap was determined by Fisher’s exact test.

### Patch-seq

We removed the cells which were labeled as “unclassified” or which belonged to rare cell types (there were less than 15 cells labeled either as “Scng” or “NP”).

For the scRNA modality we didn’t use the scVI decoder with a ZINB loss but rather just a fully connected decoder with an L2 loss on the log-normalized counts as it fitted better the data. Similarly to the smFISH/scRNA experiment, we increased the weight of the IOT loss *λ*_*IOT*_ to 0.05. Indeed our prior knowledge about connections between features across modalities consisted in connecting each gene with itself as scRNA measurements were available for both modalities. However, in contrast with the smFISH experiment where only a few dozens of genes had been measured in both modalities, here all genes could be connected, making the prior information much stronger than in previous cases. This resulted in the sinkhorn regularization not being necessary to obtain a good mixing of the two modalities, hence we set *λ*_*r*_ to 0. Moreover, the sets of cells from the two modalities being actually two independent subsets from the exact same dataset, we could expect very little heterogeneity between the cell populations present in each modality and increased the transported mass parameter from 0.5 to 0.75 in this experiment.

## Data availability

**Cell lines**. We retrieve a scCAT-seq (RNA+ATAC) dataset with 205 cells from three cancer cell lines (HCT116, HeLa-S3, K562). Data is available in the Supplementary Materials of the original publication^15^. **PBMC 10X**. We retrieve a 10X Genomics Multiome (RNA+ATAC) dataset available at https://www.10xgenomics.com/datasets/pbmc-from-a-healthy-donor-no-cell-sorting-10-k-1-standard-2-0-0. **OP Multiome and OP Cite**. We retrieve a Multiome (RNA+ATAC) and a Cite-seq bone marrow dataset from the Open Problems challenge^47^. The GEO accession number is GSE194122 and the data is available at https://www.ncbi.nlm.nih.gov/geo/query/acc.cgi?acc=GSE194122. *BMCITE*. We retrieve a CITE-seq (RNA+ADT) bone marrow dataset from Stuart et al.^27^, the GEO accession number is GSE128639 and the data is available at https://www.ncbi.nlm.nih.gov/geo/query/acc.cgi?acc=GSE128639. **Smartseq cortex**. We retrieve a scRNA-seq mouse somatosensory cortex dataset from Zeisel et al.^56^ using scvi-tools’s helper function *scvi*.*data*.*cortex*. The data is available at https://storage.googleapis.com/linnarsson-lab-www-blobs/blobs/cortex/expression_mRNA_17-Aug-2014.txt. **smFISH**. We retrieve an osmFISH mouse somatosensory cortex dataset from Codeluppi et al.^55^ using scvi-tools’s helper function scvi.data.cortex. The data is available at http://linnarssonlab.org/osmFISH/osmFISH_SScortex_mouse_all_cells.loom. **3omics RNA**. We retrieved a scRNA-seq dataset of PBMCs from a Covid study^59^ and selected cells from all healthy patients. The GEO accession number is GSE150728 and the data is available at https://www.ncbi.nlm.nih.gov/geo/query/acc.cgi?acc=GSE150728. **3omics ATAC**. We retrieve a scATAC-seq dataset of PBMCs and Bone marrow cells from an hematopoietic study in which we select the four batches of PBMCs (“PBMC_Rep1”, “PBMC_Rep2”, “PBMC_Rep3”, “PBMC_Rep4”). The GEO accession number is GSE129785 and the data is available at https://www.ncbi.nlm.nih.gov/geo/query/acc.cgi?acc=GSE129785. **3omics CyTOF**. We retrieve a CyTOF dataset of PBMCs from a Covid study in which we select an experimental batch of healthy cells (Batch B). The data is available at 10.5281/zenodo.5139560 under the name “CBD-KEY-CYTOF-WB.tar.gz”. **Patch neurons**. We retrieve a Patch-seq dataset of mouse primary motor cortex cells^68^. The scRNA counts are available with GEO accession number GSE163764 at https://www.ncbi.nlm.nih.gov/geo/query/acc.cgi?acc=GSE163764 and neuronal morphological reconstructions are available at https://download.brainimagelibrary.org/3a/88/3a88a7687ab66069/.

## Code availability

*Package*. The Python package for scConfluence is hosted at https://github.com/cantinilab/scconfluence. It can be installed easily by running pip install scconfluence. *Reproducibility*. Code to reproduce the experiments and figures is available at https://github.com/cantinilab/scc_reproducibility.

## Acknowledgements

The project leading to this manuscript has received funding from the French government under management of Agence Nationale de la Recherche as part of the “Investissements d’avenir” program, reference ANR-19-P3IA-0001 (PRAIRIE 3IA Institute). In addition this work has been funded by the Agence Nationale de la Recherche (ANR) JCJC project scMOmix, the Inception program (Investissement d’Avenir grant ANR-16-CONV-0005) and the INSERM project ITMO Cancer MIC APL-EpiNet. The work of G. Peyré was supported by the French government under management of Agence Nationale de la Recherche as part of the ‘Investissements d’avenir’ program, reference ANR19-P3IA-0001 (PRAIRIE 3IA Institute). We thank Mélanie Ridel for the administrative support. Figure elements, including the icons of modalities and datasets, were created with BioRender.com.

