## Supplementary text, tables and figures for "scConfluence : single-cell diagonal integration with regularized Inverse Optimal Transport on weakly connected features"

The Supplementary Information is organized as follows:


- Page 2 of this PDF: Supplementary Note 1
- Pages 3 to 5: Supplementary Tables 1 to 5
- Pages 6 to 13: Supplementary Figures 1 to 9

**Supplementary Note 1**

The proof only uses two classical results in OT theory. Firstly, there is a primal-dual relationship linking the optimal plan $\mathbf{Q}_\varepsilon(\mathbf{a}, \mathbf{b})$ to an optimal pair of potentials $(\mathbf{f}, \mathbf{g})$ such that:

$$\mathbf{Q}(\mathbf{a}, \mathbf{b}) = \exp(\frac{1}{\varepsilon}(\mathbf{f} \oplus \mathbf{g} - \mathbf{c}_{L_2}(\mathbf{a}, \mathbf{b}))$$ (eq plan)

Secondly, the optimal value of the entropic OT problem can be expressed with this optimal pair of potentials:

$$(\frac{1}{n_1}\mathbf{1})^T\mathbf{f} + (\frac{1}{n_2}\mathbf{1})^T\mathbf{g} = OT_\varepsilon^{+\infty}(\mathbf{a}, \mathbf{b}, c_{L_2})$$ (eq marginals)

$$\begin{aligned} KL(\mathbf{P}|\mathbf{Q}_\varepsilon(\mathbf{a}, \mathbf{b})) &= \langle \mathbf{P}, \log(\frac{\mathbf{P}}{\mathbf{Q}_\varepsilon(\mathbf{a}, \mathbf{b})})\rangle \\ &= \langle \mathbf{P}, \log(\mathbf{P})\rangle - \langle \mathbf{P}, \log(\mathbf{Q}_\varepsilon(\mathbf{a}, \mathbf{b}))\rangle \\ &= \langle \mathbf{P}, \log(\mathbf{P})\rangle - \langle \mathbf{P}, \frac{1}{\varepsilon}(\mathbf{f} \oplus \mathbf{g} - \mathbf{c}_{L_2}(\mathbf{a}, \mathbf{b}))\rangle \\ &= \langle \mathbf{P}, \log(\mathbf{P})\rangle + \frac{1}{\varepsilon}\langle \mathbf{P}, \mathbf{c}_{L_2}(\mathbf{a}, \mathbf{b}))\rangle - \frac{1}{\varepsilon}\langle \mathbf{P}, \mathbf{f} \oplus \mathbf{g}\rangle \end{aligned}$$ (Eq. Proof calcul 1)

$$\begin{aligned} \langle \mathbf{P}, \mathbf{f} \oplus \mathbf{g}\rangle &= (\mathbf{P}\mathbf{1})^T\mathbf{f} + (\mathbf{P}^T\mathbf{1})^T\mathbf{g} \\ &= (\frac{1}{n_1}\mathbf{1})^T\mathbf{f} + (\frac{1}{n_2}\mathbf{1})^T\mathbf{g} \\ &= OT_\varepsilon^{+\infty}(\mathbf{a}, \mathbf{b}, c_{L_2}) \end{aligned}$$ (Eq. Proof calcul 2)

**Supplementary Table 1.** List of the datasets used in this paper and their characteristics.

| Dataset Name | Technology | Modalities | Organism | Tissue | Cells (after QC) | Labels | Batches | Reference |
|---|---|---|---|---|---|---|---|---|
| Cell lines | scCAT-seq | RNA, ATAC | Human | Cell lines | 206 | 3 | 1 | Liu, L. et al[1] |
| PBMC 10X | 10X Multiome | RNA, ATAC | Human | PBMC | 9,378 | 14 | 1 | 10X genomics |
| OP Multiome | 10X Multiome | RNA, ATAC | Human | Bone marrow | 69,249 | 22 | 13 | Luecken, M et al.[2] |
| BMCITE | Cite-seq | RNA, ADT | Human | Bone marrow | 30,672 | 27 | 1 | Stuart, T. et al.[3] |
| OP Cite | Cite-seq | RNA, ADT | Human | Bone marrow | 90,261 | 31 | 12 | Luecken, M et al.[2] |
| Smartseq cortex | Smartseq2 | RNA | Mouse | Somato sensory cortex | 3005 | 6 | 1 | Zeisel, A. et al.[4] |
| smFISH | osmFISH | RNA | Mouse | Somato sensory cortex | 4530 | 6 | 1 | Codeluppi, C. et al.[5] |
| 3omics RNA | Seq-Well | RNA | Human | PBMC | 16627 | 12 | 6 | Wilk, A. J. et al.[6] |
| 3omics ATAC | 10X scATAC-seq | ATAC | Human | PBMC | 21261 | 18 | 4 | Satpathy, A. T. et al.[7] |
| 3omics CyTOF | Helios CyTOF | Protein | Human | PBMC | 43232 | 21 | 1 | Covid-19 Multi-omics Blood Atlas Consortium[8] |
| Patch neurons | Patch-seq | RNA, morphologies | Mouse | Primary motor cortex | 1214 | 7 | 1 | Scala, F. et al[9] |

**Supplementary table 2.** List of genes identified as differentially expressed in both the scRNA and scATAC gene activities for each B cell cluster.

| Cluster 0 | Cluster 1 | Cluster 2 |
|---|---|---|
| AIM2, BLK, CCDC50, CD1C, COTL1, CTSH, HLA-DPB1, HLA-DQA1, KCNN4, MAP4K1, MARCKS, MS4A1, OAZ1, POU2AF1, POU2F2, PPP1R15A, PTPN1, RALGPS2, SCIMP, SCRN1, SPIB, SYK, SYNGR2, TBC1D9, TFEC, TLR10, TNFRSF13B, UBC, UBE2J1, WDFY4 | BCL7A, BTG1, BTLA, DGKD, FAM129C, FOXP1, HLA-DMB, ICOSLG, PCDH9, TCL1A, TSPAN13, YBX3 | ACTG1, ANXA1, AOAH, APMAP, APOL3, APOL6, ARF1, ARHGEF1, ARL4C, ATP1A1, ATP2B4, ATP8B2, BCL11B, BIN2, BTN3A1, BTN3A2, BTN3A3, C11orf21, C1orf21, CAB39, CALM1, CALR, CAMK4, CANX, CBLB, CCDC88C, CCL5, CCND2, CCND3, CCSER2, CD2, CD247, CD300A, CD3D, CD3E, CD48, CD5, CD6, CD63, CD8A, CD96, CDC42SE2, CEP78, CFL1, CFLAR, CHD3, CST7, CTBP2, CTSW, CX3CR1, CYLD, CYTH1, CYTIP, DGKA, DGKZ, DIAPH1, DIP2A, DOK2, DYNC1H1, DYRK2, EFHD2, EIF3A, ESYT2, EVL, F2R, FBXW5, FCGR3A, FGD3, FGFBP2, FKBP5, FLNA, FOSL2, FYN, GBP5, GIMAP1, GIMAP4, GIMAP5, GIMAP7, GLG1, GNLY, GUK1, GZMA, GZMB, GZMH, HELZ, HERC1, HIPK1, HLA-A, HLA-B, HLA-C, HLA-E, HLA-F, HSPA5, HSPA8, ID2, IFITM1, IFITM2, IGF2R, IL10RA, IL2RB, IL32, IL6ST, IL7R, INPP4A, IQGAP2, ITGA6, ITGAL, ITGB1, ITGB2, ITK, ITM2B, JADE2, KCNAB2, KDM3A, KIAA1551, KLF13, KLRB1, KLRD1, KLRF1, KPNB1, LASP1, LCK, LCP1, LCP2, LDHB, LEF1, LINC00861, LITAF, LPIN2, MAN1A1, MATK, MBP, MCL1, MGAT4A, MLLT6, MSN, MYBL1, MYH9, MYL12A, MYL12B, MYL6, MYO1F, NCAM1, NDFIP1, NKG7, PAG1, PARP8, PBXIP1, PCED1B-AS1, PDE3B, PDZD4, PFN1, PIK3IP1, PIK3R1, PIM1, PIP4K2A, PPP2R5C, PREX1, PRF1, PRKCQ, PRMT2, PRPF38B, PRSS23, PTPN12, PTPRA, RAB27A, RAP1B, RARRES3, RASA3, RASAL3, RASGRP1, RASSF1, RASSF5, RBL2, RBMS1, REST, RICTOR, RNF125, RNF213, RPS6KA3, RUNX3, S100A4, S100A6, S1PR5, SAMD3, SAMD9, SELPLG, SEMA4D, SH2D1A, SIGIRR, SKP1, SLAMF7, SLC9A3R1, SLFN5, SMARCA2, SORL1, SPN, SPOCK2, SPTAN1, SRGN, SSBP3, ST3GAL1, STAT1, STK10, STK38, SYNE1, SYNE2, SYTL2, TAX1BP1, TBX21, TCF7, TES, TESPA1, TMC8, TNFAIP3, TNFRSF1B, TNIK, TPP2, TPST2, TRABD2A, TRAF3IP3, TRANK1, TSHZ1, TXK, UBE2G2, UTRN, VIM, WDR82, WIPF1, ZAP70, ZFP36L2, ZNF91 |

**Supplementary table 3.** List of genes identified as differentially expressed in both the scRNA and scATAC gene activities for each monocyte cluster.

| Cluster 4 | Cluster 5 |
|---|---|
| ACTN1, ADAM15, ADAM8, AGTRAP, AHNAK, AHR, ALDH2, ANXA1, ANXA6, APLP2, APOL3, APP, ARHGAP26, ARHGEF40, ASGR2, ATP6V0B, ATP6V1A, ATP6V1F, BAZ2B, BHLHE40, BLVRB, BST1, CAPG, CASP4, CCDC149, CCDC88A, CCR1, CCR2, CD14, CD163, CD1D, CD63, CD84, CD93, CDA, CIITA, CKAP4, CLEC4A, CLEC4E, CLMN, CMIP, CMTM3, COMT, CPD, CPM, CREG1, CRISPLD2, CRTAP, CSF3R, CST3, CTSA, CTSH, CTSS, CXCR4, CXXC5, CYFIP1, CYP1B1, CYP27A1, DGKD, DHRS4, DYSF, EFHD2, EIF4G3, EMB, F13A1, F5, FAM129A, FAM198B, FCN1, FERMT3, FES, FKBP5, FLOT1, FNDC3B, FPR1, FRMD4B, G0S2, GAPDH, GLRX, GLT1D1, GM2A, GPX1, GRN, H2AFY, HEBP2, HEXB, HK2, HLA-A, HLA-DQB1, HLA-DRA, HLA-DRB5, HPSE, HSD17B4, IDH1, IGF2R, IGSF6, IL13RA1, IL4R, IL6R, IL6ST, IMPA2, IQGAP2, IRF2BP2, IRS2, ITGA5, ITGAM, ITGB2, IVNS1ABP, KCTD20, KDM4B, KDM7A, KIAA0040, KIF13A, KLF10, LAMP2, LAMTOR1, LAPTM5, LAT2, LBR, LGALS2, LGALS3, LINC00963, LITAF, LPGAT1, LRP1, LTB4R, LTBR, LY86, LYZ, MAPK14, MARC1, MARCO, METTL9, MGST1, MID1IP1, MLKL, MLXIP, MNDA, MPEG1, MSRB1, MYCL, MYO1F, NCF4, NFE2, NFKBIA, NLRP12, NLRP3, NR4A2, NRGN, OSCAR, P4HB, PADI2, PADI4, PARP8, PEA15, PER1, PID1, PKM, PLA2G7, PLBD1, PLD3, PLEKHO1, PLXND1, PPIF, PRRC2B, PSTPIP1, PTPRE, PYGL, QPCT, QSOX1, RAB11FIP1, RAB27A, RAB3D, RBM47, RBP7, RIT1, RPS16, RPS8, S100A10, S100A12, S100A6, S100A8, S100A9, S1PR3, SCPEP1, SEMA4D, SEPT2, SGK1, SIRPA, SLC2A3, SLC40A1, SMARCD3, SOCS3, SORL1, STAB1, STX3, SULF2, SYK, TAGLN2, TALDO1, TAPBP, TBC1D9, TET3, THBS1, TLR4, TMEM173, TMEM205, TMEM71, TNFAIP2, TNFAIP3, TNFRSF1A, TPP1, TPT1, TREM1, TRIB1, TRPS1, TSPO, TXN, VEGFA, VIM, WLS, XRN2, YBX3, YWHAE, ZNF385A, ZNF467 | ACOT9, ALDH3B1, ARRDC2, ASAH1, C15orf39, CDKN1B, CDKN1C, CKB, CRIP1, CSF1R, CSK, CYTH1, FAM110A, FAM49A, FCGR3A, FOXO1, FZD1, GNAI2, GPI, GPR137B, HES4, HLA-E, HSBP1, HSPA8, IFITM2, IQSEC1, KLF11, KLF12, KLF2, KLF7, KNDC1, LFNG, LRMP, LRRC25, LRRFIP1, LYL1, LYN, MAFB, MEG3, METRNL, MGLL, MRPS35, MS4A7, MTSS1, MYO1G, MYOF, NECAP2, PDPK1, PIK3CG, PILRA, PKN1, PPP1R17, PSAP, PTP4A2, PTPN1, PTPRC, RALB, RHOB, RIN3, RNH1, RRAS, SAT1, SFT2D2, SLC44A2, SMAD2, SNX5, SPG11, SPRED1, SSBP4, TPTEP1, UNC119, VPS35, WARS, WAS, XIAP, YBX1, ZBTB7A, ZFAND5, ZFR, ZNF703 |

**Supplementary Table 4.** Results of the differential expression analysis for the cluster 6 of monocytes in the tri-omics experiment. The output of scanpy's rank_gene_groups method is displayed for two known marker genes of monocyte-derived dendritic cells.

| names | scores | logfoldchanges | pvals | pvals_adj |
|---|---|---|---|---|
| CCR7 | 2.67e+01 | 5.40e+00 | 2.06e-157 | 9.74e-154 |
| CD2 | 2.15e+01 | 5.61e+00 | 2.75e-102 | 8.79e-100 |

**Supplementary Table 5.** List of the number of features measured in each modality of the datasets used and the number of neurons in hidden layers of the autoencoders .

| Dataset Name | Number of features | Hidden size |
|---|---|---|
| Cell lines/RNA | >10000 | 64 |
| Cell lines/ATAC | >10000 | 64 |
| PBMC 10X/RNA | >10000 | 64 |
| PBMC 10X/ATAC | >10000 | 64 |
| OP Multiome/RNA | >10000 | 64 |
| OP Multiome/ATAC | >10000 | 64 |
| BMCITE/RNA | >10000 | 64 |
| BMCITE/ADT | 25 | 20 |
| OP Cite/RNA | >10000 | 64 |
| OP Cite/ADT | 134 | 32 |
| Smartseq cortex | >10000 | 64 |
| smFISH | 33 | 25 |
| Covid RNA | >10000 | 64 |
| Hemato ATAC | >10000 | 64 |
| Covid CyTOF | 48 | 32 |
| Neurons/RNA | >10000 | 64 |
| Neurons/Images | 2048 | 128 |

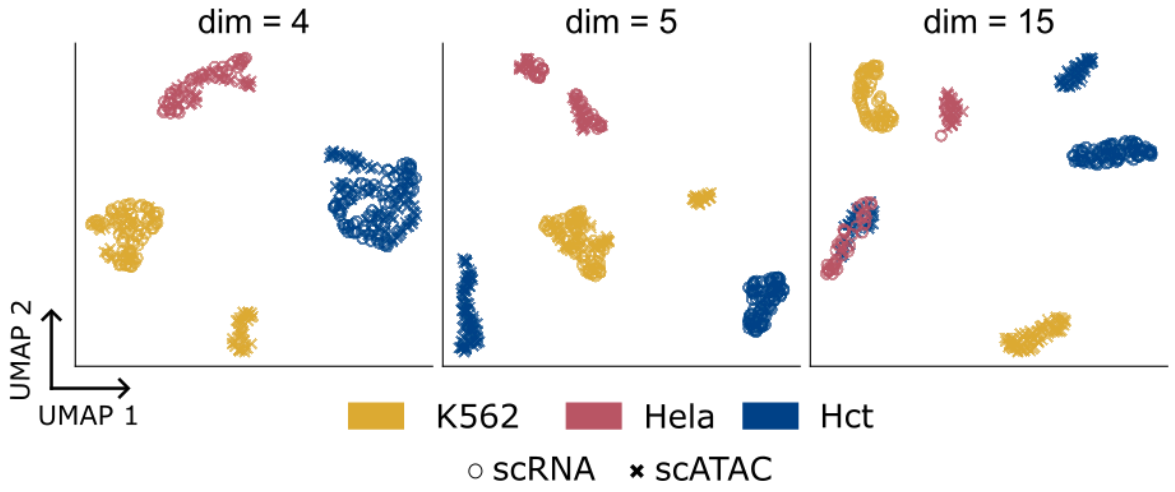


**Supplementary Figure 1.** 2D UMAP visualizations of the cell embeddings obtained by LIGER on the cell lines dataset for different values of the dimension of the latent space. Different colors in these UMAP plots correspond to the three different cell lines present in the data while the shape of the point markers correspond to the modality of origin of each cell (scRNA, scATAC).

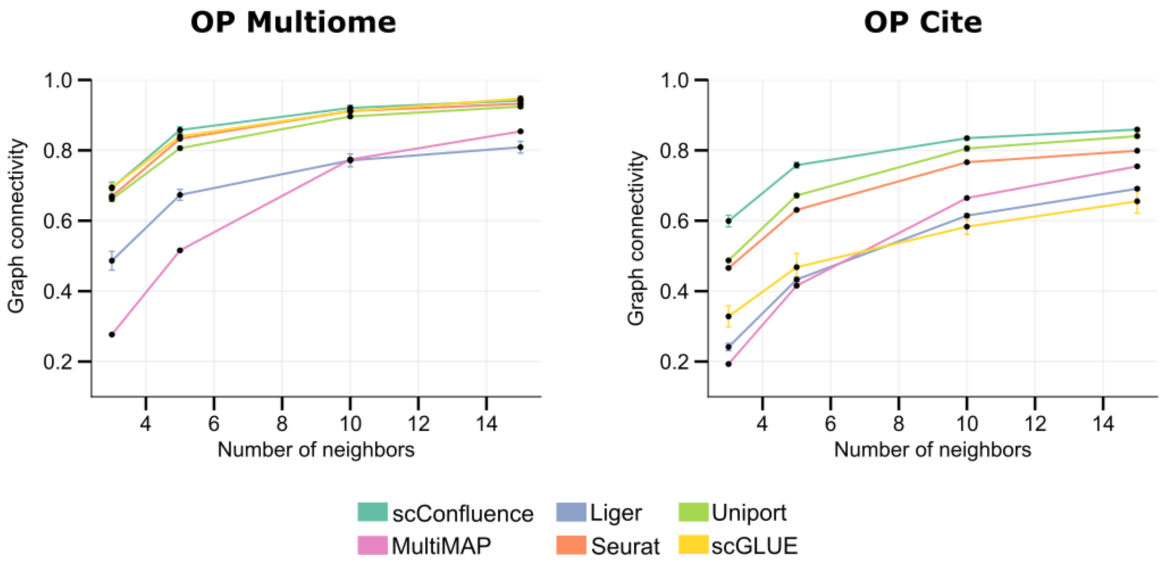


**Supplementary Figure 2.** Graph connectivity scores for the six benchmarked methods (scConfluence, Seurat, Liger, MultiMAP, Uniport and scGLUE) in the two bone marrow datasets which contained multiple experimental batches.

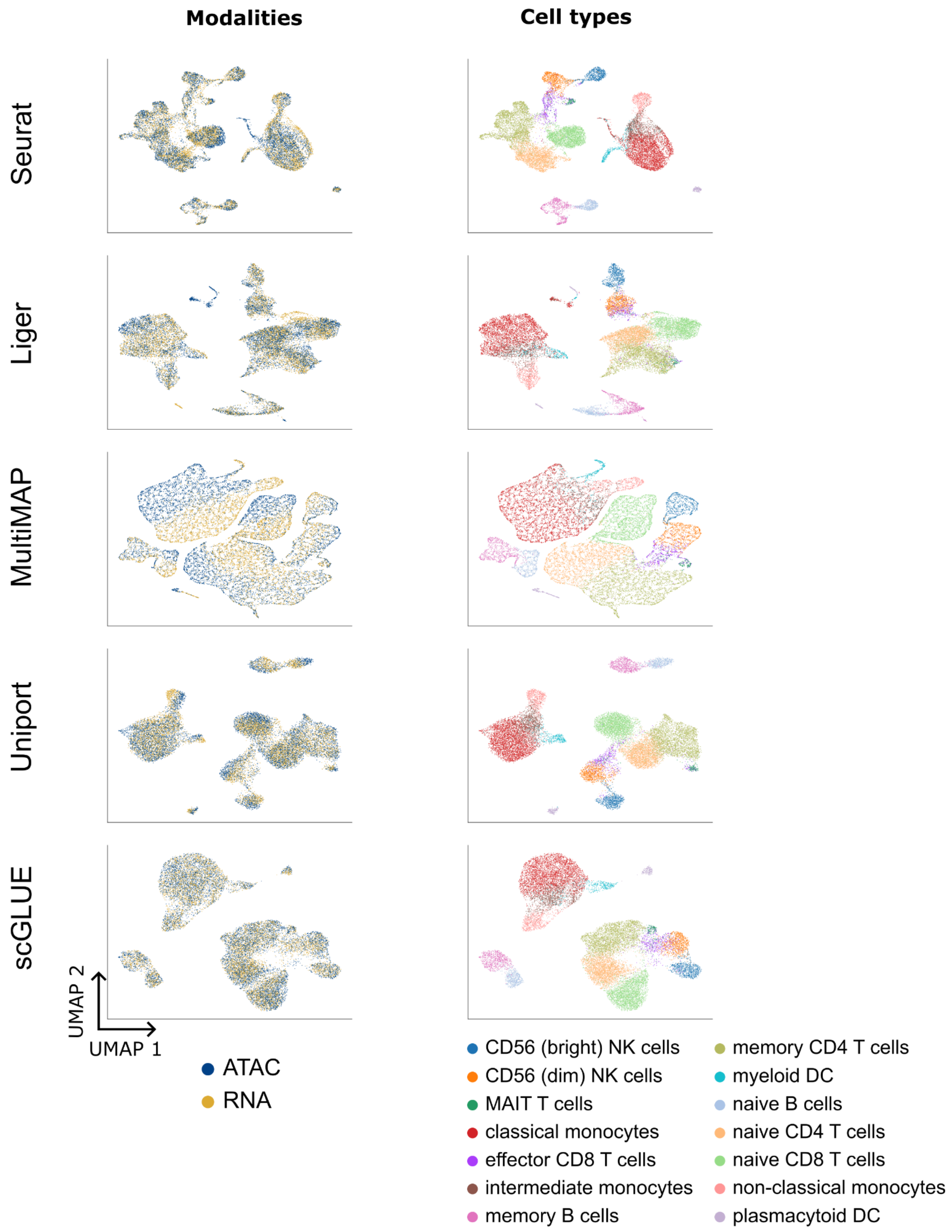


**Supplementary Figure 3.** 2D UMAP visualizations of the cell embeddings obtained by the five baselines (Seurat, Liger, MultiMAP, Uniport and scGLUE) on the *PBMC 10X* dataset. Cells are colored based on their modality of origin and their cell type annotation.

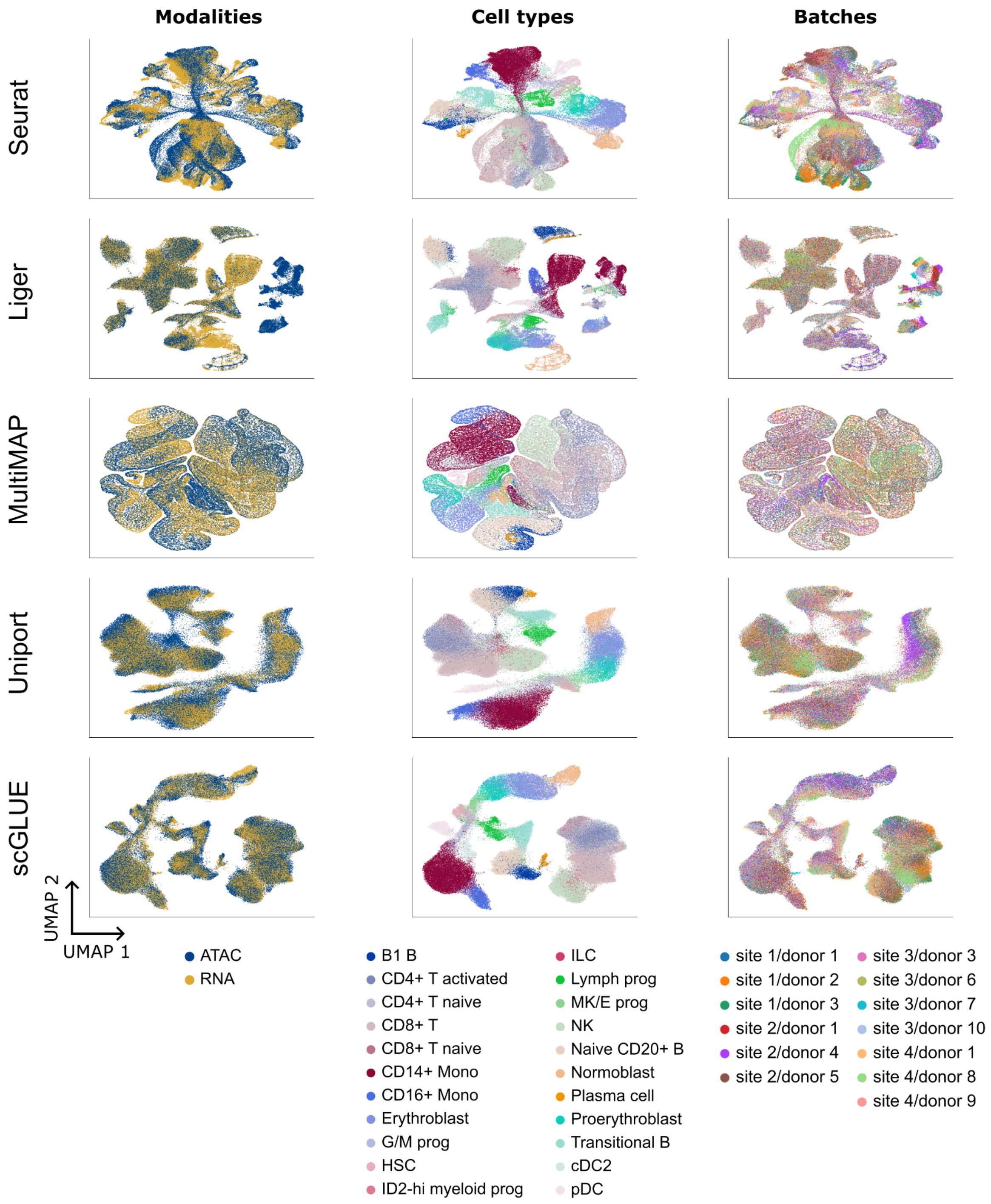


**Supplementary Figure 4.** 2D UMAP visualizations of the cell embeddings obtained by the five baselines (Seurat, Liger, MultiMAP, Uniport and scGLUE) on the *OP Multiome* dataset. Cells are colored based on their modality of origin, their cell type annotation or their batch of origin.

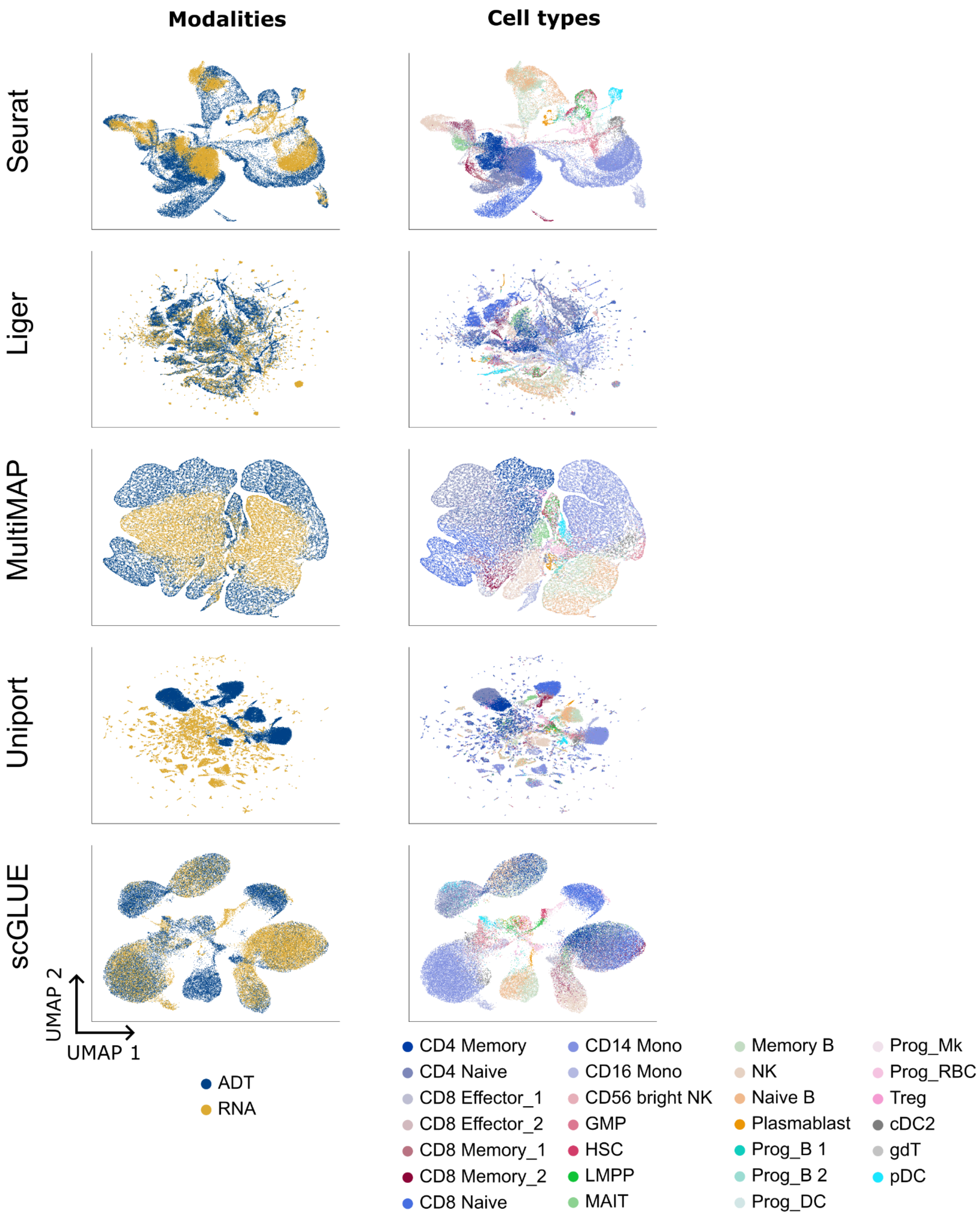


**Supplementary Figure 5.** 2D UMAP visualizations of the cell embeddings obtained by the five baselines (Seurat, Liger, MultiMAP, Uniport and scGLUE) on the *BMCITE* dataset. Cells are colored based on their modality of origin or their cell type annotation.

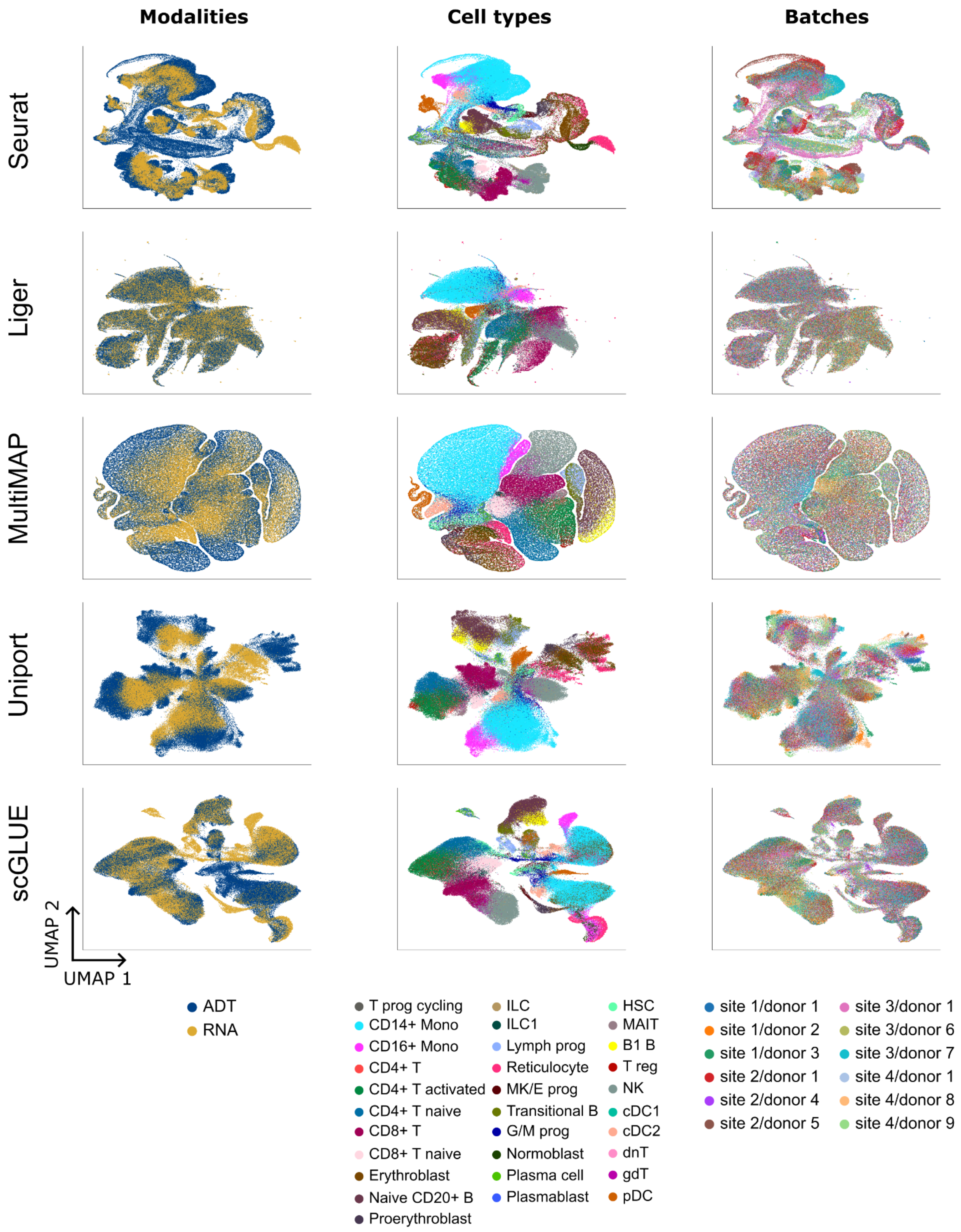


**Supplementary Figure 6.** 2D UMAP visualizations of the cell embeddings obtained by the five baselines (Seurat, Liger, MultiMAP, Uniport and scGLUE) on the *OP Cite* dataset. Cells are colored based on their modality of origin, their cell type annotation or their batch of origin.

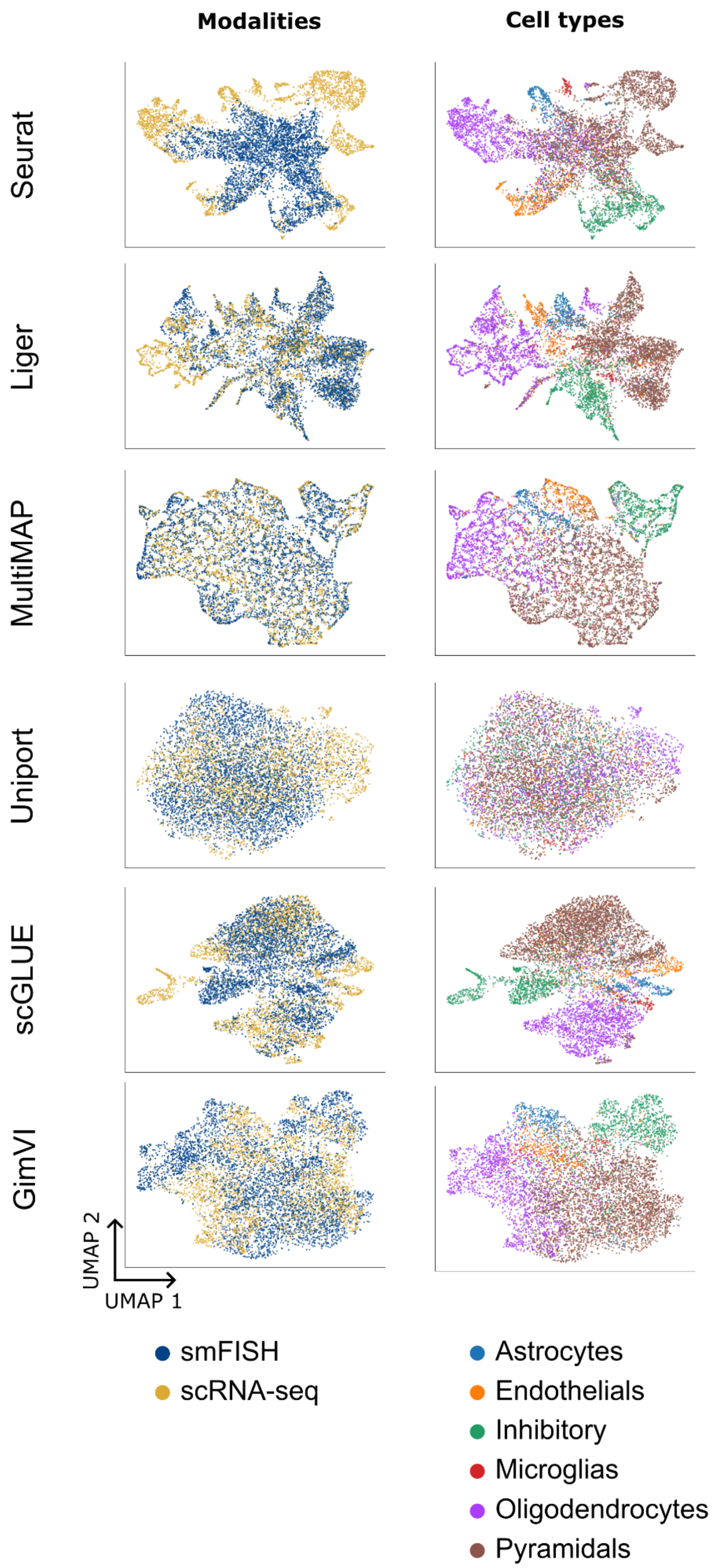


**Supplementary Figure 7.** 2D UMAP visualizations of the cell embeddings obtained by the six baselines (Seurat, Liger, MultiMAP, Uniport, scGLUE and GimVI) on the *scRNA/smFISH* dataset. Cells are colored based on their modality of origin and their cell type annotation.

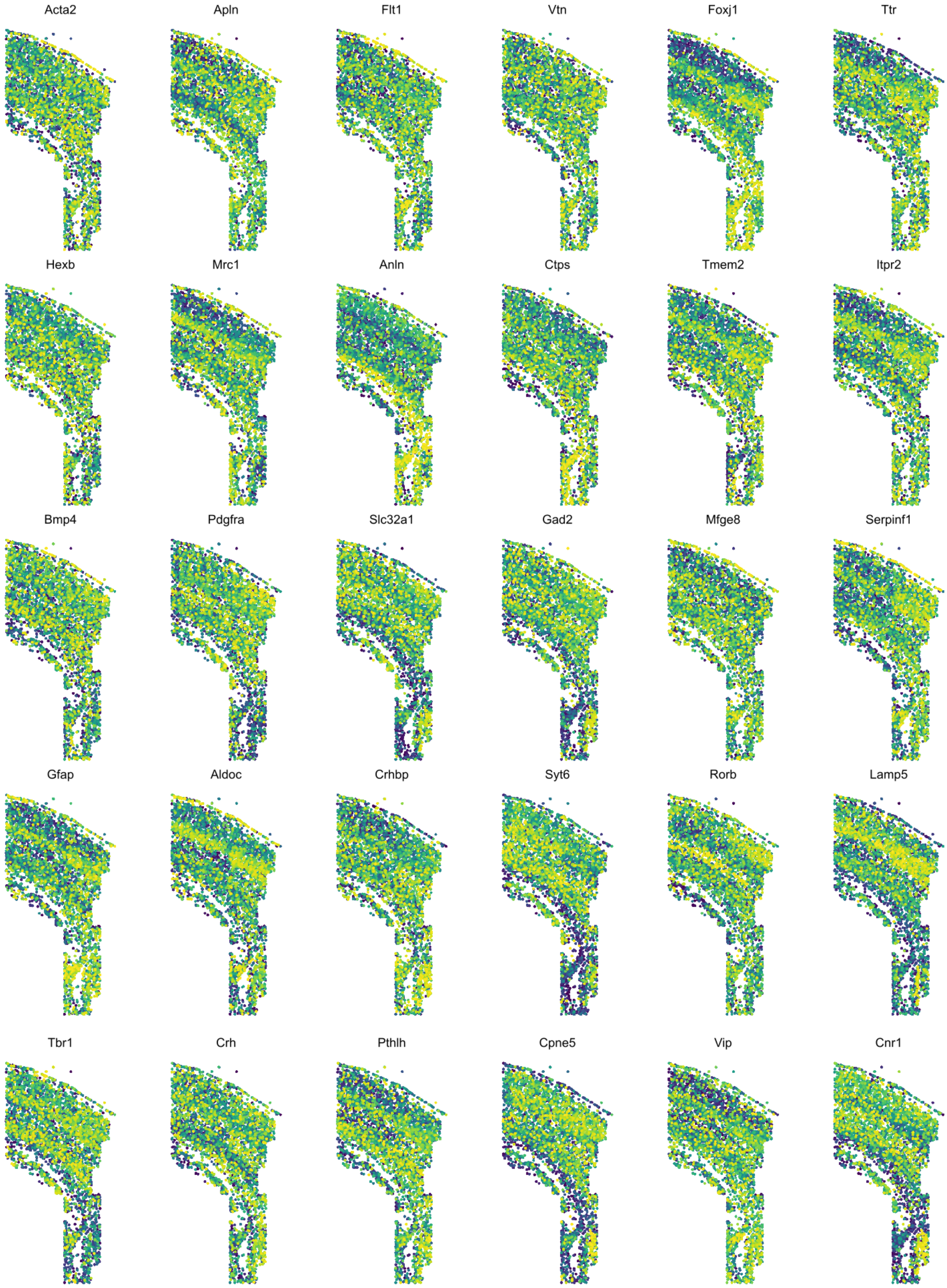


**Supplementary Figure 8.** Spatial pattern of expression of scConfluence's imputations on the thirty held-out smFISH genes which were not displayed in Figure 6.

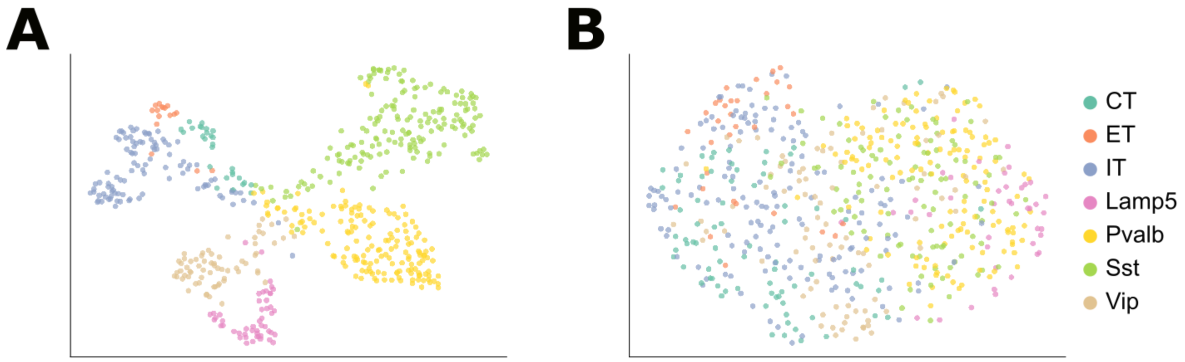


**Supplementary Figure 9. Unimodal embeddings of scRNA-seq and morphologies before integration.** UMAP visualizations of cell embeddings obtained by training independent autoencoders on the two modalities,  (a) scRNA counts and (b) neuronal morphologies, without integrating them together. Cell embeddings are colored by their transcriptomic cell type annotations.
